# MG2Act: A Mechanism-Inspired Sequential Attention Framework for Molecular Glue Degradation Prediction

**DOI:** 10.64898/2026.08.08.739980

**Authors:** Zhiyao Zhuang, Dan Teng, Xiaojing Xu, Senbiao Fang, Yifan Wang, Menglu Hou, Ling Ge, Shanwen Yuan, Miaoyuan Yang, Lei Cheng, Zongqun Zhang, Qi He, Zhuoyue Li, Ximing Xu, Shumin Ma, Sulin Zhang, Xiao Wang, Mingyue Zheng, Chong Qin

## Abstract

Molecular glue degraders act by inducing productive proximity between an E3 ligase and a substrate protein. For most characterized degradative glues, a small molecule first engages the E3, conditions its substrate-recognition surface, and only then enables recruitment of a compatible neo-substrate. This directionality is rarely encoded explicitly in computational models, which typically fuse molecule, E3 and target representations simultaneously. We present MG2Act, a structure-independent framework that translates this two-step logic into sequential cross-attention, using CRBN-mediated degradation as the most data-rich representative system. Starting from a curated continuous-valued benchmark of 1,207 pairs across 47 targets, a refined subset of 1,159 pairs was selected to train MG2Act after excluding rare targets. On identical processed data, MG2Act consistently outperforms machine learning baselines, robustly generalizes under strict redundancy-filtering, and responds coherently to mechanism-based perturbations. Prospective screening and zero-shot target-conditioned prioritization identified nanomolar degraders of IKZF1, CK1α and CDK4, including the non-classical IMiD-core CDK4 degrader SWC-202.

## Introduction

Targeted protein degradation expands the druggable proteome by reprogramming endogenous E3 ubiquitin ligases to eliminate disease-relevant proteins^1,2^. Within this paradigm, molecular glue degraders (MGDs) are monovalent small molecules that create or stabilize a productive interface between an E3 ligase and a substrate protein^3–6^, distinct from bifunctional PROTACs that rely on both E3-binding and substrate-binding ligands tethered by a linker^7,8^. In most characterized degradative glues, the ligand first engages an E3 ligase, most prominently cereblon (CRBN), and induces or stabilizes a neomorphic surface that enables cooperative recruitment of a non-native substrate^3–6^. CRBN-recruiting MGDs are the dominant and best-characterized class, with established activity against challenging targets including IKZF1/3^6^, CK1α^9^, GSPT1^10^, WIZ^11^, WEE1^12^, NEK7^13^ and VAV1^13^. Other mechanisms exist: cyclin K degraders such as CR8 bind the ATP pocket of CDK12-cyclin K and recruit the DDB1-CUL4-RBX1 core, illustrating a target-bound route^14^. Despite this diversity, the central design challenge is shared. MGD activity is not governed by a predefined binding geometry, but by the cooperative formation of a transient ternary interface whose productive state depends on the order and compatibility of molecular events^4,15–18^.

MGD discovery has therefore relied heavily on empirical screening and iterative optimization. Computer-aided drug design and deep learning offer a path toward scalable prioritization^19^, and protein sequence embeddings, graph neural networks and generative models have been used to predict activity or optimize linker architectures for PROTACs^20–24^. AI methods dedicated to molecular glue degraders, however, remain scarce; to our knowledge, no specialized framework has yet been reported for quantitative, target-conditioned prediction of MGD activity. Three obstacles constrain progress. First, ternary-complex modelling is impeded by induced-fit dynamics and transient interfacial contacts that current co-folding tools cannot reliably capture^18^. Second, MGD pharmacological data are sparse and heterogeneous, frequently mixing qualitative Western blot readouts with incomplete DC_50_ and D_max_ measurements. Third, existing AI frameworks for targeted degradation rely on ligand-centric representations or fuse molecule, E3 and target information in a largely simultaneous manner, without explicitly encoding the directed temporal order of molecular glue action.

This simultaneous-fusion assumption is mathematically convenient but obscures an important biological asymmetry. For most E3-first glues, ligand binding to the E3 precedes and enables the surface state that recruits a substrate. Structural studies of CRBN show this directly: ligand binding causes the surface remodeling that exposes the neomorphic interface^4–6^; only the ligand-conditioned surface is competent to recruit a neo-substrate. The process is directed in time, typically from ligand to E3 and then to substrate, and a model intended to learn molecular glue compatibility should be able to represent that direction. A model that pools E3 and target representations symmetrically may still capture statistical associations, but cannot represent the temporal asymmetry that defines many molecular glue mechanisms, leaving open the concern that apparent predictive success derives partly from ligand-activity associations propagated through redundant training examples^25^.

Encoding domain knowledge as an architectural inductive bias is a well-established strategy. Convolutional networks operationalize spatial locality, equivariant networks encode physical symmetries, physics-informed neural networks embed governing differential equations into the training objective^26^, and biology-informed networks such as P-NET represent hierarchical pathway structure directly in their connectivity^27^. We apply a related idea to directional molecular glue degradation. Using CRBN-mediated MGDs as the most data-rich representative system, MG2Act encodes the common E3-first sequence of ligand engagement, E3 surface conditioning and substrate recruitment as a two-stage attention architecture. The architecture does not prove the mechanism. Rather, the known biological order of events is represented as the model’s information flow and then evaluated through falsification-oriented perturbations.

Building on our previously published MolGlueDB^28^, the first comprehensive database dedicated to molecular glue degraders, we curated a high-quality CRBN-focused dataset as a representative benchmark for E3-first molecular glue modelling. Expert-guided standardization reconciled disparate experimental readouts into a unified continuous degradation activity score that integrates potency (DC_50_) and maximal efficacy (D_max_) through a penalty-weighted framework. The continuous-valued benchmark (1207 high-confidence compound-target pairs across 47 neo-substrates) preserves graded activity information that binary labels discard. We then built MG2Act, a structure-independent framework whose main architectural choices implement this mechanism-inspired inductive bias. Graph-based molecular representations with explicit functional-group-aware attention encode chemical priors known to govern E3 engagement; ESM-C^29^ embeddings encode the evolutionary and structural context of the E3 ligase and the recruited target; and a sequential cross-attention engine fuses these modalities in a biologically motivated order. Stage-reversal ablation, glutarimide perturbation and target swapping provide complementary tests of whether the learned signal is consistent with the expected biophysics, and prospective discovery against IKZF1, CK1α and CDK4 examines whether the signal can prioritize experimentally active degraders, including the sub-nanomolar CK1α degrader **YFC-403** and the CDK4 degrader **SWC-202** identified via hierarchical scaffold prioritization.

## Results

### Sequential cross-attention encodes E3-first molecular glue logic

Many degradative molecular glues act through a directional sequence in which a small molecule first conditions an E3 ligase surface and thereby enables recruitment of a compatible substrate. CRBN-mediated MGDs provide the best-characterized example (Figure 1a). A monovalent ligand engages the CRBN tri-tryptophan pocket and allosterically remodels its surface, generating a neomorphic interface that subsequently enables cooperative recruitment of a compatible neo-substrate^5,13^. The process is directed: target recruitment is conditional on prior ligand-E3 engagement and cannot occur in the reverse order. MG2Act is designed so that information flows through the model in this same direction (Figure 1b).

**Figure 1.**
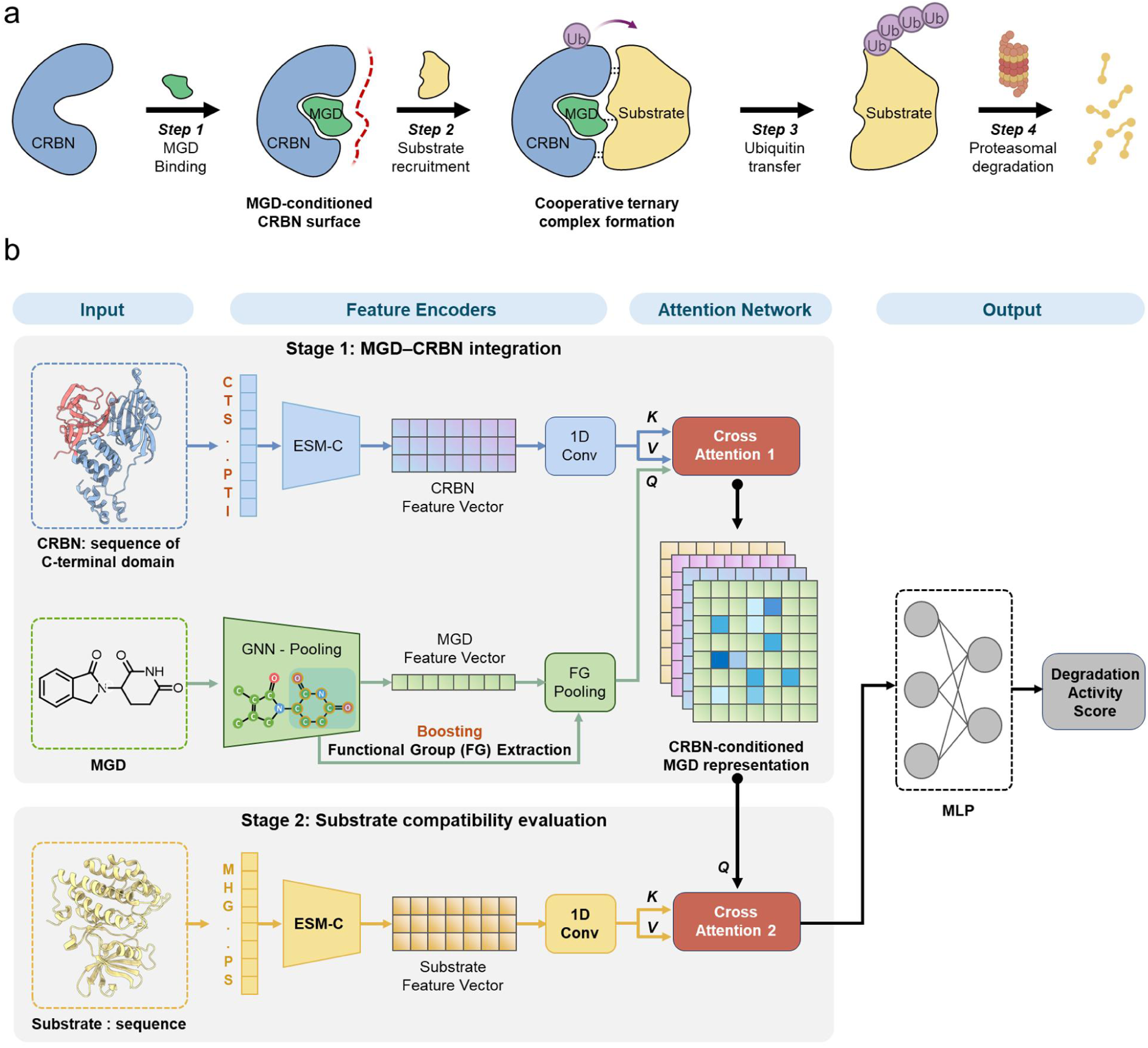
Biological mechanism of molecular glue mediated protein degradation and the architecture of the MG2Act deep learning framework. **a**, Directed E3-first molecular glue degradation, illustrated using CRBN-mediated MGDs as a representative system. Ligand engagement remodels the E3 surface, which then recruits the neo-substrate to form a productive ternary complex. **b**, Schematic of the MG2Act architecture. The two cross-attention stages (ligand → CRBN, then CRBN-conditioned ligand → target) are aligned with the two biological stages depicted in a.

Three modality-specific encoders feed the directed engine. Small molecules are represented as molecular graphs constructed with RDKit, where atoms (with elemental, hybridization and formal-charge features) are nodes and bonds are edges, and a three-layer Graph Attention Network (GAT) produces atom-level embeddings. To inject chemical priors that govern CRBN engagement, MG2Act explicitly identifies 14 functional groups per ligand, curated by filtering an initial set of 29 chemically common groups for at least 5% occurrence and merging chemically equivalent variants; each group is represented as a pooled embedding of its constituent atoms. To precisely capture the relevant biophysical interactions while minimizing evolutionary noise, we specifically utilized the C-terminal thalidomide-binding domain (CTD, residues 318–442) of CRBN rather than its full-length sequence, as the CTD harbors the tri-tryptophan pocket responsible for ligand anchoring and subsequent neo-substrate recruitment . CRBN CTD and the target protein are independently embedded by the ESM-C protein language model (300 M parameter variant) and projected into a shared latent space^27^.

The directional core of MG2Act is its two-stage cross-attention engine. In stage 1, functional-group queries from the molecule attend to CRBN residues (keys and values), producing CRBN-conditioned molecular representations that operationalize the ligand-induced surface remodeling event. In stage 2, these CRBN-conditioned representations attend to the target protein, producing target-conditioned functional-group features that operationalize cooperative neo-substrate recruitment. A functional-group boosting strategy softly amplifies the attention weights of chemically validated MGD scaffolds (boost factor λ = 2), incorporating weak chemical priors while preserving data-driven attention. The fused representation is concatenated with pooled global embeddings of CRBN and the target, and passed through a multilayer perceptron to output a continuous degradation activity score in [0, 1].

Reversing the stage order, target before CRBN, corresponds to the opposite of the known biological sequence, and we show below that it reduces model performance accordingly. The architecture therefore encodes a mechanistic hypothesis rather than adding model complexity.

### A continuous-valued activity score that couples potency and maximal degradation

Faithful modelling of molecular glue degradation requires activity annotations that capture both potency and maximal efficacy. The half-maximal degradation concentration DC_50_ describes the concentration at which 50% of the achievable target degradation has occurred, while the maximal degradation D_max_ is the upper plateau of the dose response and reflects how completely the target can be removed. These two parameters are pharmacologically distinct. A compound can show an attractive DC_50_ yet leave a substantial fraction of the target intact, in which case downstream pathway suppression remains incomplete and therapeutic benefit is limited. Target proteins often must be depleted below a threshold before a cellular phenotype is reversed, and partial degradation can translate into partial functional rescue at best^13,30,31^. This issue is general to molecular glue degraders, because glue-induced ternary complexes can produce potent but incomplete degradation depending on substrate geometry, cell context and ubiquitination efficiency. An activity score that does not penalize incomplete D_max_ risks rewarding compounds that appear potent in dose response but cannot achieve sufficient target clearance for therapeutic effect.

Public MGD data are heterogeneous, comprising disparate DC_50_ values, single-point Western blot readouts and qualitative activity ranges. Traditional pipelines collapse this heterogeneity into binary labels, discarding graded structure-activity information essential for lead optimization.

From MolGlueDB^28^, we curated a CRBN-focused subset (Figure 2a) through literature re-examination and expert-guided standardization. Reported DC_50_ values were mapped to a normalized potency score (Score_DC50_) on a 0–1 scale (>10 µM → 0; 1–10 µM → 0.2; 0.1–1 µM → 0.4; 10–100 nM → 0.6; 1–10 nM → 0.8; <1 nM → 1.0). When available, maximal degradation was incorporated as a penalty term (Penalty_Dmax_: >80% → 0; 60–80% → 0.33; 40–60% → 0.67; <40% → 1.0). The final continuous degradation activity score is

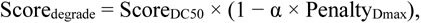

where α is a tunable weight controlling the D_max_ contribution. The multiplicative form penalizes molecules with adequate potency but partial efficacy, reflecting the biological observation that effective therapeutic degradation typically requires substantial target clearance^30,31^. For entries lacking D_max_ annotations, the penalty contribution was minimized to preserve robustness to incomplete reporting.

**Figure 2.**
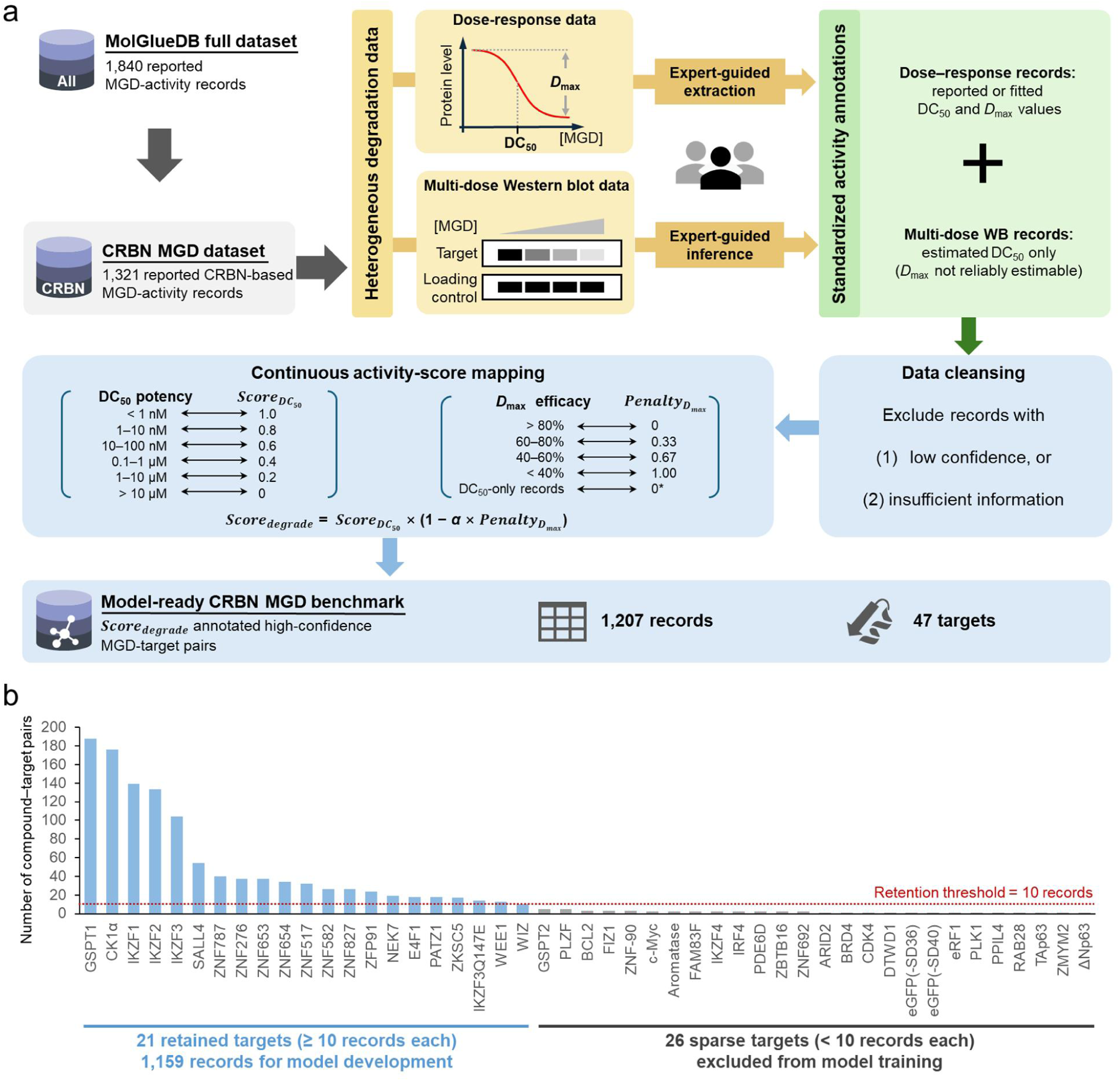
Construction and content of the continuous-valued CRBN molecular glue activity dataset. **a**, Dataset construction pipeline and assignment of continuous scores from graded DC_50_ potency segments and D_max_-derived penalties. **b,** Total number of targets and entries in the degradation dataset. **c**, Targets and activity distribution of the curated dataset; the lower panel shows the distribution before and after incorporation of the D_max_ penalty.

After automated and manual quality control, the curated dataset initially comprised 1,207 high-confidence compound-target pairs across 47 targets (Figure 2b). To ensure adequate per-target statistics and mitigate potential target-specific learning biases arising from data sparsity, we excluded rare targets with fewer than 10 associated entries. This filtering process yielded a refined final subset of 1,159 pairs across 21 targets. This threshold prevents the model from overfitting to underrepresented biological contexts or memorizing isolated ligand-activity shortcuts, while ensuring that each target can be robustly represented across all data splits. Consequently, the dataset was split 8:1:1 into training, validation, and test sets, adjusted to maintain a relatively uniform distribution of targets across subsets. The continuous scoring preserves graded activity information and explicitly couples potency and maximal degradation, providing a principled benchmark for regression-based modelling and virtual screening.

### MG2Act outperforms machine learning baselines and improves early enrichment

Trained under a strict train, validation and test split, MG2Act achieves R² = 0.5950 and RMSE = 0.1751 on the independent test set (Table 2, Figure 3b), indicating reliable ranking of degradation activity across diverse molecular glue and target pairs. Table 1 summarizes the final hyperparameters, and Table 2 reports the validation-set sweep used to select them.

**Figure 3.**
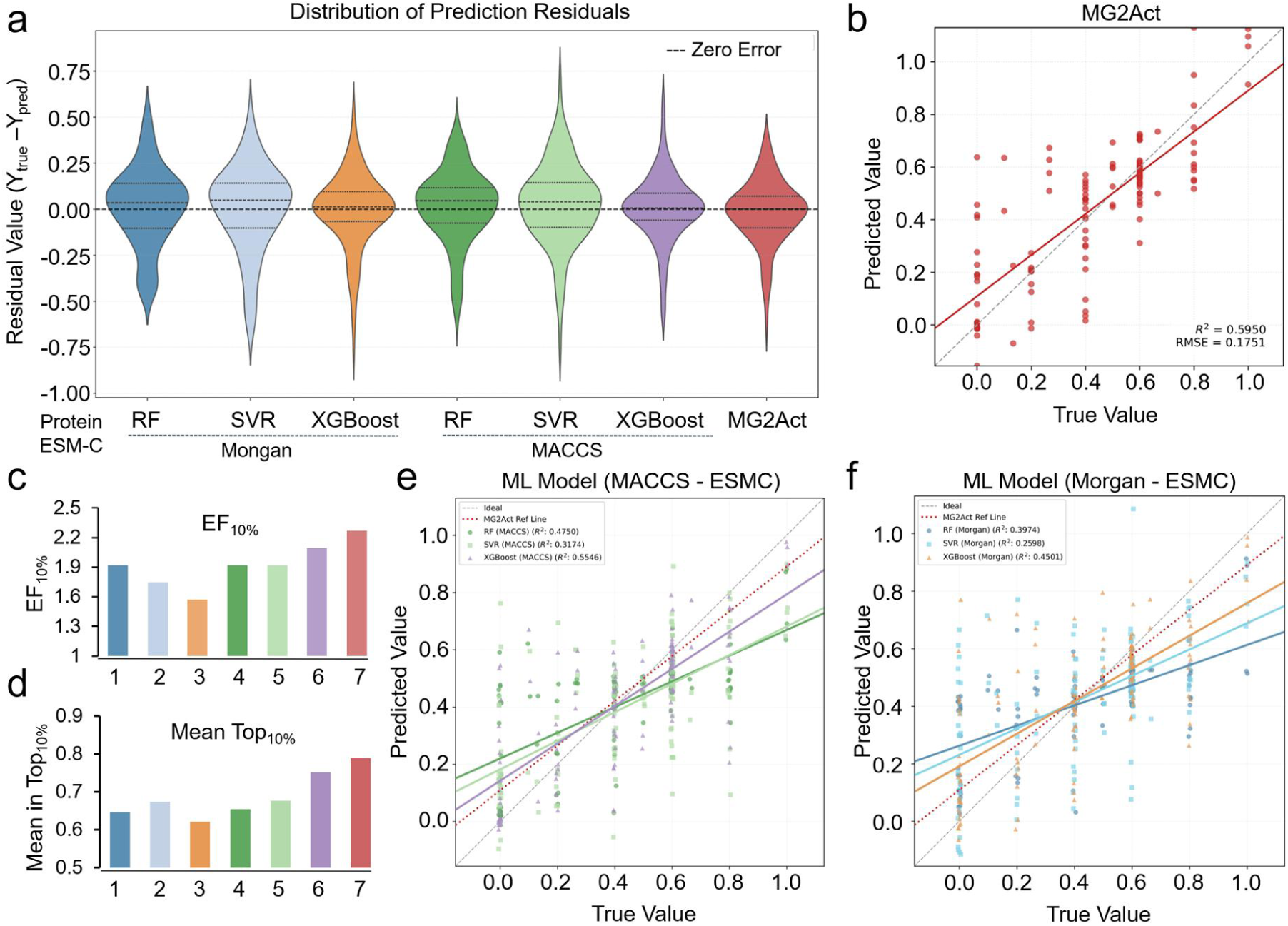
MG2Act outperforms baseline models. Baselines are 1, RF (Morgan-ESMC); 2, SVR (Morgan-ESMC); 3, XGBoost (Morgan-ESMC); 4, RF (MACCS-ESMC); 5, SVR (MACCS-ESMC); 6, XGBoost (MACCS-ESMC); 7, MG2Act, **a**, Violin plots of prediction residuals (observed − predicted). Dashed lines indicate the median and interquartile range; the horizontal dashed line marks zero error. **b**, Scatter of MG2Act predictions versus experimental values; grey dashed line, ideal y = x; red line, fitted regression. **c**, Enrichment factor at the top 10% (EF_10%_). **d**, Mean activity among the top 10% ranked compounds across molecular representations. **e, f**, Predicted versus observed values for RF, SVR and XGBoost using MACCS fingerprints (points perturbed by 0.05 horizontally for visualization). For all models compared, protein sequences are encoded using ESMC embeddings, identical to the representation used in MG2Act; all baselines therefore take both protein (ESMC) and ligand features as input.

**Table 1.** Dimensions and hyperparameters of the MG2Act model.

| Module | Parameter description | Value / Type |
| --- | --- | --- |
| Protein | Protein sequence feature extractor | ESM-C 300M |
|  | Dimensionality reduction | Conv |
|  | Embedding dimension | 64 |
| Molecule | Graph neural network type | GAT |
|  | Number of GNN layers | 3 |
|  | Hidden feature dimension | 128 |
|  | Initial atom feature dimension | 44 |
|  | Initial bond feature dimension | 10 |
|  | Functional-group aggregation | Max pooling |
| | FG boosting factor $\lambda$ | 2 |
| **Fusion** | Functional-group attention enabled | True |
|  | Attention heads | 4 |
|  | Transformer decoder layers | 2 |
| **MLP** | Fully connected layer 1 / layer 2 | 128 / 64 |
| **Training** | Max epochs / batch size | 200 / 8 |
| | Optimizer / learning rate | Adam / $1 \times 10^{-4}$ |
|  | Early-stopping patience / dropout | 30 / 0.1 |

**Table 2.** Validation-set hyperparameter optimization.^a^.

| Category | Setting | MAE | RMSE |
| --- | --- | --- | --- |
| **Learning rate** | $3 \times 10^{-6}$ | 0.1858 | 0.2436 |
| | $1 \times 10^{-5}$ | 0.1465 | 0.2086 |
| | $3 \times 10^{-5}$ | 0.1467 | 0.1987 |
|  | **$1 \times 10^{-4}$** | **0.1322** | **0.1885** |
| | $3 \times 10^{-4}$ | 0.1459 | 0.1969 |
| **Batch size** | 1 | 0.2279 | 0.2561 |
|  | 2 | 0.1281 | 0.1893 |
|  | 4 | 0.1375 | 0.1903 |
|  | **8** | **0.1322** | **0.1885** |
|  | 16 | 0.1404 | 0.1890 |
|  | 32 | 0.1401 | 0.1932 |
|  | 64 | 0.1370 | 0.1932 |
|  | 128 | 0.1532 | 0.2087 |
| **Protein projection** | Linear | 0.1332 | 0.1918 |
|  | MLP | 0.1494 | 0.1978 |
|  | **Convolution** | **0.1322** | **0.1885** |
| **E3 representation** | Full E3 sequence | 0.1552 | 0.2005 |
|  | **E3 functional domain** | **0.1322** | **0.1885** |
| **Molecular encoder** | **GAT** | **0.1322** | **0.1885** |
|  | GNN | 0.1413 | 0.1926 |
| **Hidden dimension** | 64 | 0.1419 | 0.1994 |
|  | **128** | **0.1322** | **0.1885** |
| **Hidden layers** | 2 | 0.1390 | 0.1928 |
|  | **3** | **0.1322** | **0.1885** |
| **Attention heads<sup>b</sup>** | 2 | 0.1321 | 0.1914 |
|  | **4** | **0.1322** | **0.1885** |
|  | 8 | 0.1332 | 0.1867 |

<sup>a</sup>Bold marks the configuration adopted in the final model. <sup>b</sup>4 attention heads was selected over 8, as the latter offered no substantial performance gains while unnecessarily inflating model complexity.

To benchmark MG2Act against conventional multimodal regression models, we trained Random Forest (RF), Support Vector Regression (SVR) and XGBoost using statically concatenated representations of the molecular glue, CRBN and target protein. Specifically, each baseline combined a Morgan or MACCS molecular fingerprint with pooled ESM-C embeddings of the CRBN C-terminal thalidomide-binding domain and the corresponding target protein. The strongest baseline, XGBoost with MACCS and protein embeddings, achieved R^2^ = 0.5550, compared with R^2^ = 0.5950 for MG2Act. Although these baselines access the same three information sources, they represent each modality as a fixed global feature vector and do not explicitly model residue-level interactions or the ordered ligand–CRBN–target conditioning process. This comparison therefore establishes the overall advantage of MG2Act over conventional tri-modal feature aggregation, whereas the contribution of sequential cross-attention and E3-first information flow was further isolated through the architecture-matched concatenation and stage-reversal ablations described below. The validation-set sweep (Table 2) shows that MG2Act’s molecular and protein encoders are each well matched to the task: replacing the GAT molecular encoder with a vanilla GNN raises RMSE from 0.1885 to 0.1926, and restricting the E3 input to the CRBN functional domain rather than the full sequence improves RMSE from 0.2005 to 0.1885, consistent with the model focusing on the region biophysically responsible for ligand engagement.

Under screening-relevant conditions, in which compounds with degradation scores at least 0.5 are defined as highly active, MG2Act achieves the highest early enrichment among all methods. The enrichment factor at the top 10% (EF_10%_) is 2.27, and the mean true activity among the top 10% ranked compounds is the largest of any method (Figure 3c, d). In practical screening, only a small fraction of computationally prioritized candidates is synthesized, and MG2Act preferentially places potent degraders in that fraction.

### Reversing the directional stage order reduces performance

If MG2Act’s predictive success arises in part from representing the directed mechanism (ligand → CRBN → target), then forcing the model to process information in the reverse order should impair performance, whereas a model that merely learned ligand activity associations would be less affected by reversing the protein conditioning order. We therefore performed an ablation in which the order of the two cross-attention stages was reversed (target first, then CRBN), keeping every other architectural element fixed. The full ablation suite is reported in Table 3.

**Table 3.** Ablation studies probing the mechanistic content of MG2Act.

| Category | Configuration | MAE | RMSE | R<sup>2</sup> |
| --- | --- | --- | --- | --- |
| **Full MG2Act model** | fusion: **sequential attention**<br>protein representation: **ESM-C**<br>FG boost factor: $\lambda = 2$ | **0.1239** | **0.1751** | **0.5950** |
| **Fusion variant** | reversal | 0.1447 | 0.1978 | 0.4833 |
|  | concatenation | 0.1483 | 0.1991 | 0.4765 |
| **Representation variant** | ACC | 0.1336 | 0.1810 | 0.5673 |
|  | one-hot | 0.1373 | 0.1877 | 0.5348 |
| **Functional-group boost variant** | $\lambda = 1.5$ | 0.1352 | 0.1885 | 0.5306 |
| | $\lambda = 1$ (disabled) | 0.1485 | 0.1977 | 0.4840 |

Reversing the stage order causes R² to drop from 0.595 to 0.4833 while the parameter count, encoders and training data remain unchanged. The model is therefore sensitive to the property that distinguishes the known biology from a symmetric interaction: the direction of information flow contributes to the predictive signal, not merely the presence of protein conditioning. Concatenation (no cross-attention) yields R² = 0.4765, indicating that the structured cross-attention and its biologically motivated order each contribute independently to performance.

Two further ablations probe complementary aspects of the architecture. Replacing pretrained ESM-C embeddings with handcrafted protein features (amino-acid composition (ACC) and one-hot encoding) degrades performance even when the sequential attention is preserved (R² = 0.5673 and 0.5348, respectively), indicating that evolutionary-scale protein representations supply residue-level structural context that the directed attention exploits.

Functional-group boosting shows a monotonic improvement as λ increases from 1 (off) to 2 (R² 0.4840 → 0.5306 → 0.5950), reflecting the benefit of a moderate chemical prior that emphasizes pharmacologically relevant motifs without overriding the data-driven attention.

Each architectural commitment in MG2Act—the directional order of cross-attention, the evolutionary protein embeddings and the chemical-prior-informed molecular encoder—therefore contributes to the mechanism-inspired design, and removing or reversing any one of them reduces performance.

### A de-redundancy benchmark separates compatibility learning from memorization

A characteristic feature of the molecular glue field is that a single CRBN-binding scaffold can recruit several distinct neo-substrates. Lenalidomide is the canonical example: it engages CRBN through its glutarimide handle and, in different cellular contexts, drives the degradation of IKZF1 and IKZF3 in multiple myeloma^32,33^ and CK1α in del(5q) myelodysplastic syndrome^34^. Thalidomide and pomalidomide show overlapping but distinct neo-substrate spectra^6^. This polypharmacology arises directly from the underlying mechanism: the same ligand can remodel the CRBN surface in a manner compatible with more than one G-loop or G-loop-like degron motif^13^. As a result, a given SMILES routinely appears multiple times in any MGD activity table, each time paired with a different target and activity value. A model can therefore exploit a shortcut: if the same molecular feature appears repeatedly at similar activity levels in training, an apparent protein-conditioned performance can be recovered in part by memorizing ligand-to-score associations and ignoring the target. Similar concerns have been raised for graph-based protein-ligand affinity models^25^ and apply with particular force to molecular glue datasets in which the same compound is intentionally tested across many neo-substrates.

We therefore constructed a stringent de-redundancy benchmark. From the initial test set, every entry whose SMILES *and* activity score were both replicated in the training set was removed, while molecules associated with distinct targets or distinct activity values were retained. The resulting de-redundant test set comprises 79 unique molecule-target pairs whose exact (SMILES, score) pairs were never seen during training.

Under this benchmark, the fingerprint baselines collapse to R² ≤ 0.2710 (**SI Figure 2**), confirming that a substantial share of their apparent performance on the standard test set was driven by label-redundancy memorization. MG2Act, by contrast, retains R² = 0.4223. When the easy path of memorization is removed, baseline models lose most of their signal, whereas a model that has learned protein-conditioned compatibility continues to generalize.

### Mechanism-based perturbations move predictions in the expected direction

Ablation and de-redundancy tests probe the model’s average behavior. To examine whether individual predictions are aligned with biophysical expectations, we performed two complementary perturbation analyses on the independent test set. Each perturbation is designed so that biology predicts a specific, sign-known shift in the model output. These tests do not establish causality, but provide falsification-oriented evidence that the model is not relying solely on ligand-level correlations.

#### Perturbation 1: Glutarimide N-methylation

The glutarimide moiety is the chemical handle by which CRBN-engaging MGDs anchor into the CRBN binding pocket through a conserved hydrogen-bond pattern^5,6^. *N*-methylation of the imide nitrogen introduces steric hindrance and disrupts this hydrogen bonding, and is known experimentally to abolish CRBN engagement. The chemical change is minimal but its biological consequence is severe, so a mechanism-consistent model should respond with a sharp drop in predicted activity even though the rest of the molecule is unchanged.

Across the test set, *N*-methylation of the glutarimide drives the mean MG2Act-predicted activity from 0.4396 to 0.1036 (**SI Figure 3**), a pronounced attenuation triggered by a single chemical modification. The average predicted score decreased by 0.3360, with 84.25% of the molecules predicted to exhibit a downward trend, which aligns consistently with the underlying biological mechanism. The structural change is localized, so a ligand-memorization predictor would be expected to move less systematically. The fact that MG2Act shifts its prediction in the direction expected from CRBN binding-pocket biology is consistent with internal representations that capture CRBN engagement requirements.

#### Perturbation 2: Target swapping

If MG2Act predictions are jointly conditioned on the molecular glue and the recruited target, computationally substituting the target should shift the predicted activity in a target-coherent way. We systematically replaced the target protein of each test entry with one of four highly represented CRBN neo-substrates (CK1α, GSPT1, IKZF1 and IKZF2), excluding entries originally associated with the substituted target to avoid trivial baseline effects. The resulting paired comparisons (Figure 4a–e) show substantial, target-specific shifts in predicted activity, with distinct ΔScore distributions for each substituted target. While the shift for CK1α did not reach statistical significance, pronounced shifts were observed for GSPT1, IKZF1, and IKZF2. The same molecule does not receive the same prediction across targets.

**Figure 4.**
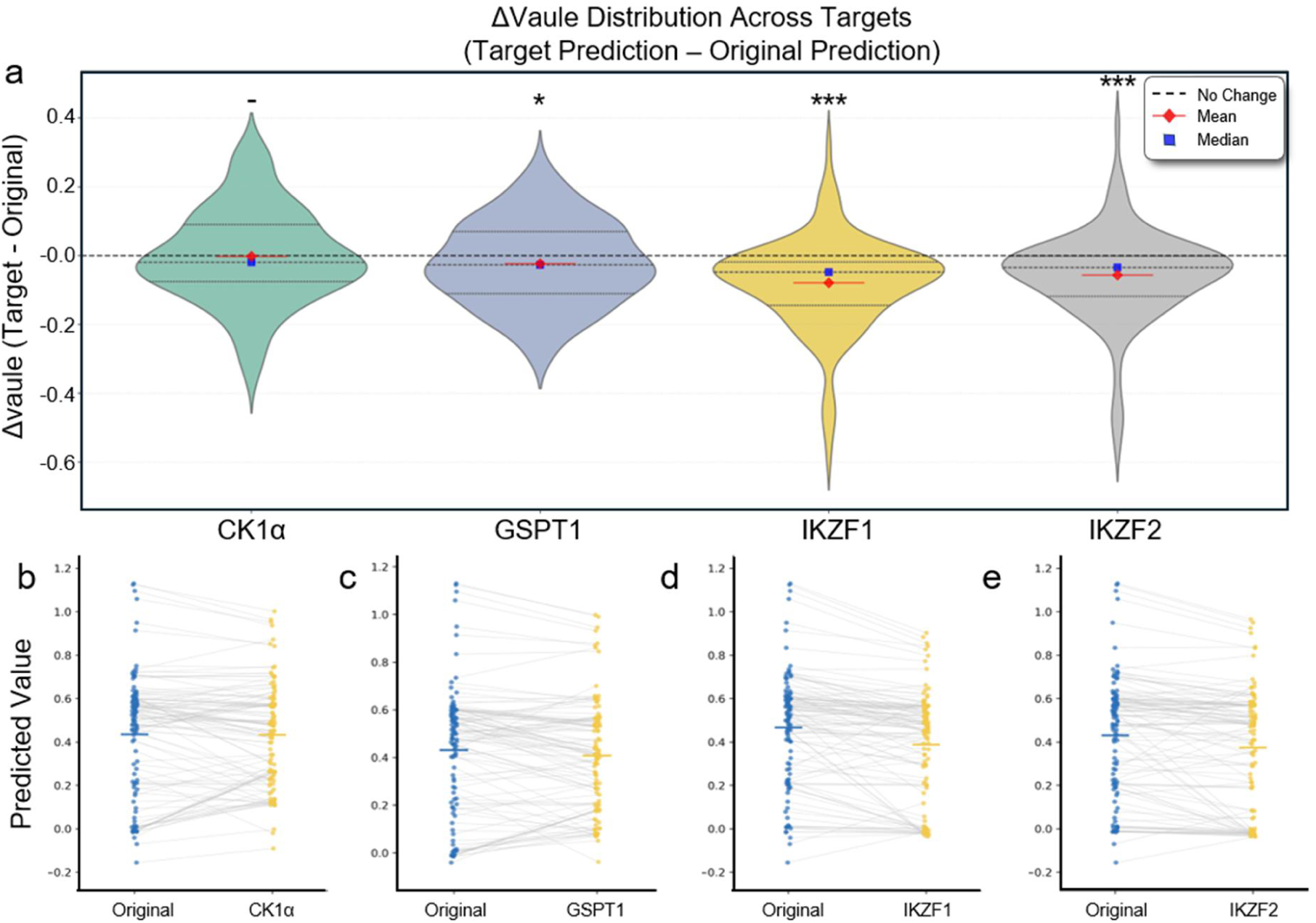
Target-specific and chemical perturbations move MG2Act predictions in the biologically expected direction. **a**, Violin plots of ΔScore distributions across substituted targets (ΔScore = target prediction − original prediction) for CK1α( n = 108, p = 0.871), GSPT1( n = 108, p < 0.05), IKZF1( n = 113, p < 0.001) and IKZF2( n = 113, p < 0.001); horizontal dashed line, no change; red diamonds, mean; blue squares, median. Statistical significance was assessed using a two-sided one-sample t-test against zero (*p < 0.05, **p < 0.01, ***p < 0.001). **b–e**, Paired comparisons of original versus target-substituted predictions for each compound (CK1α, GSPT1, IKZF1, IKZF2 respectively); each grey line connects the same compound.

These shifts cannot be explained from ligand features alone or from target-specific biases in isolation. They require the model to condition its predictions on compatibility among the molecular glue, CRBN and the recruited target, the three-way logic that MG2Act’s directed architecture is designed to capture.

Together with the stage-reversal ablation and the de-redundancy benchmark, these perturbations form a triangulation: three independent classes of intervention (architectural reversal, chemical modification and target substitution) each shift the prediction in the direction expected from the biology, consistent with a mechanism-consistent compatibility signal for CRBN engagement → surface remodeling → target recruitment, rather than ligand memorization.

Prospective virtual screening identifies nanomolar IKZF1 and CK1α degraders. Having established that MG2Act captures ordered ligand–CRBN–target dependencies, we next evaluated whether its learned signal could support prospective compound prioritization. MG2Act was applied to an in-house collection of approximately 80 CRBN-biased compounds synthesized in our previous studies and not included in MolGlueDB. These molecules retain a glutarimide or dihydrouracil moiety to support CRBN engagement while incorporating diverse substituents that may modulate neosubstrate recruitment. We focused on IKZF1 and CK1α, two canonical but mechanistically distinct CRBN neosubstrates. Although both proteins are recruited through the CRBN–molecular-glue interface, their distinct degron geometries and ligand-dependent degradation profiles provide a stringent test of target-conditioned prioritization.^32–36^

For each target, compounds were prospectively selected using a predefined score-stratified and scaffold-diversity-aware procedure. The ranked library was divided into high-(top 40%), intermediate-(40%–70%) and low-scoring (70%–100%) tiers. Following scaffold-based clustering within each tier, eight, six and four structurally representative compounds, respectively, were selected for each target. Pooling the two independently generated candidate sets yielded 28 unique compounds. The target-conditioned predictions produced partially overlapping but distinct ranking profiles, suggesting contributions from both shared CRBN-directed chemical features and neosubstrate-dependent recruitment compatibility. .

The 18 candidates selected for each target were experimentally evaluated for cellular degradation by Western blotting, followed by dose–response characterization of prioritized active compounds. MG2Act scores showed moderate and consistently positive correlations with the experimentally derived degradation score for both CK1α (Pearson r=0.595) and IKZF1 (r=0.660; Figure **5a**). More importantly for prospective screening, active compounds were preferentially enriched in the high-scoring tiers. Four of the eight compounds sampled from the top 40% for each target met the prespecified high-activity criterion. Relative to the activity prevalence within the corresponding prospectively sampled panels, this translated into enrichment factors of 1.67 for IKZF1 and 2.50 for CK1α. Thus, although continuous-score agreement remained moderate, MG2Act provided practically useful early enrichment of active degraders.

**Figure 5.**
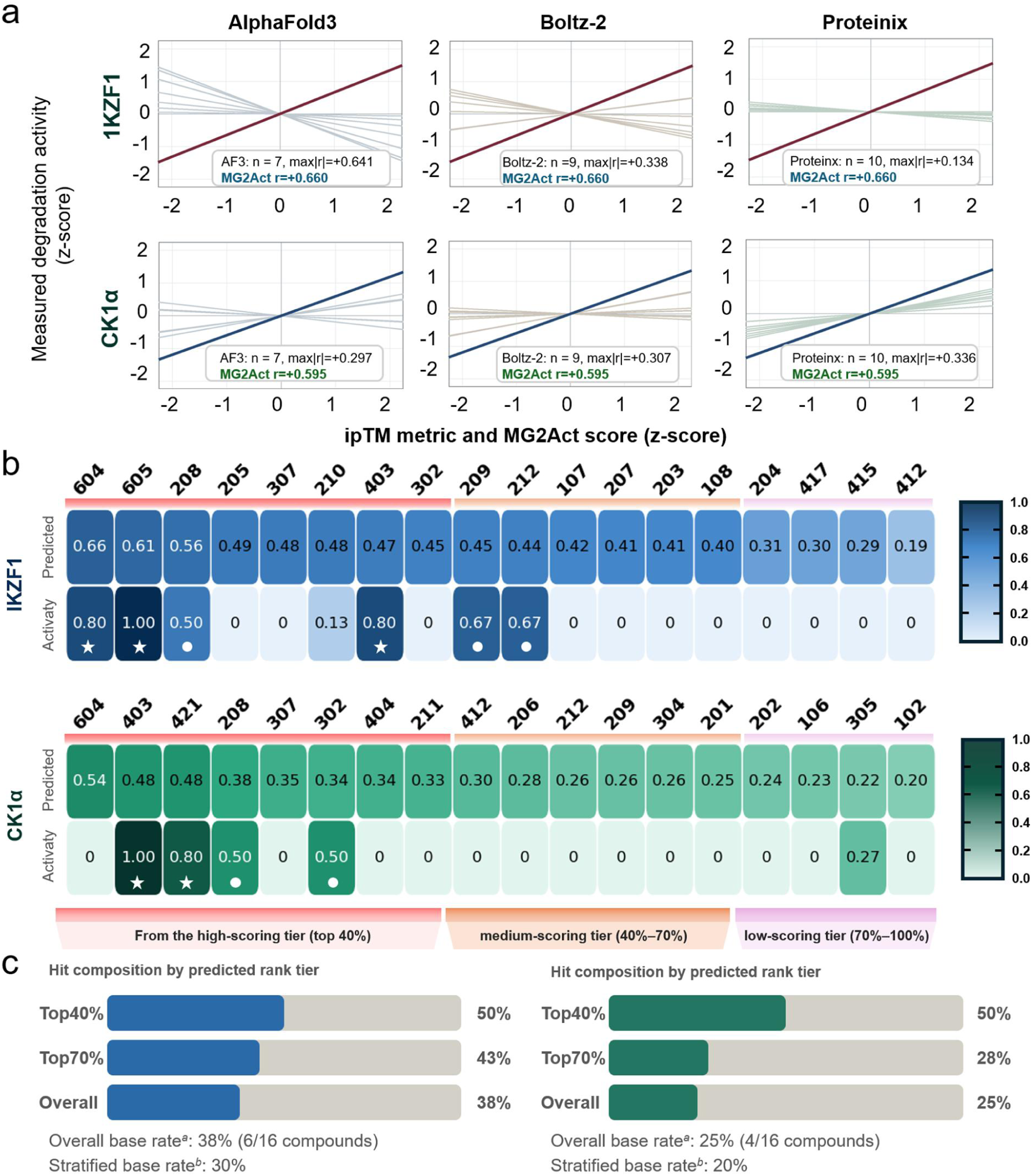
Prospective prioritization of IKZF1 and CK1α molecular glue degraders. **a**, Correlations of measured degradation activity with MG2Act scores and co-folding-derived ipTM metrics for CK1α (top) and IKZF1 (bottom). Light lines show linear fits for individual ipTM metrics from AlphaFold3 (n = 7), Boltz-2 (n = 9) and Protenix (n = 10), whereas dark lines show MG2Act. Variables were z-score normalized within each target; line slopes therefore equal Pearson r. Insets report the MG2Act r and the ipTM metric with the largest |r| **b**, Predicted and measured activities of the prospectively selected compounds, ordered by MG2Act score and grouped into high-(pink), medium-(orange), and low-scoring(lilac) tiers. White symbols indicate highly active compounds. **c**, Proportions of highly active compounds across predicted rank tiers and the full validation set. ^a^Overall base rate: The unadjusted empirical hit rate calculated directly from the total tested sample, defined as the total number of identified active compounds divided by the total number of sampled compounds (n = 18). ^b^Stratified base rate: The population-weighted hit rate estimated to correct for sampling bias introduced by tier-stratified sampling. It is calculated as ∑(*p_i_* × *w_i_*), where *p_i_* is the observed hit proportion within score tier ***i***, and ***w****_i_* is the true population proportion of tier ***i*** in the entire library (Top 40% tier = 0.40; Medium 40%–70% tier = 0.30; Low 70%–100% tier = 0.30).

To assess whether co-folding-derived interface-confidence metrics could provide comparable activity-ranking signals, we examined the associations between experimentally measured degradation activity and prespecified ipTM-related metrics generated by AlphaFold3^37^, Boltz-2^38^ and ProteniX^39^. MG2Act scores showed moderate and consistently positive correlations with experimental activity for both CK1α (r=0.595) and IKZF1 (r=0.660). In contrast, the magnitude and direction of the correlations for ipTM-related metrics varied across co-folding methods and target systems, with no consistently transferable relationship observed across both neosubstrates. These results suggest that ipTM metrics may capture target-specific interface-confidence information, whereas MG2Act provides a more consistent activity-aligned signal across the evaluated systems. The ipTM-based comparison is shown in Figure 5a, and the complete exploratory analysis of all model-reported metrics is provided in Table S1Crucially for prospective applications, MG2Act demonstrated marked early enrichment of highly active degraders across both target contexts (Figure **5c**). To account for tier-stratified sampling across the chemical library, we calculated a weighted stratified base rate to estimate the overall population prevalence of active compounds (Methods). For IKZF1, candidate prioritization within the top 40% score tier yielded a hit rate of 50%, representing a substantial enhancement over the stratified library base rate of 30%. Similarly, for CK1α, the hit composition within the top 40% tier reached 50%, more than doubling the stratified library base rate of 20%. These findings confirm that MG2Act score-stratified sampling effectively enriches active molecular glue degraders, offering significant practical efficiency for prospective screening.

Prospective validation identified several potent degraders among the prioritized candidates. YFC-605 received a high MG2Act score for IKZF1 (0.614) and induced nanomolar IKZF1 degradation, with a DC_50_ of 0.67 nM and a D_max_ of 83.10% (Figure **6a**). YFC-421 was prioritized for CK1α with a predicted score of 0.482 and showed robust cellular degradation (DC_50_ = 7.60 nM, D_max_=86.74%). YFC-403 ranked within the high-scoring region for both target contexts and exhibited potent degradation of IKZF1 (DC_50_=5.92nM, D_max_ = 82.66%) and CK1α (DC_50_=0.35 nM, D_max_=100%). YFC-403 was therefore the most potent CK1α degrader identified in this prospective set. Collectively, these compounds demonstrate that high MG2Act rankings can translate into nanomolar cellular degradation and distinct target-dependent activity profiles.

**Figure 6.**
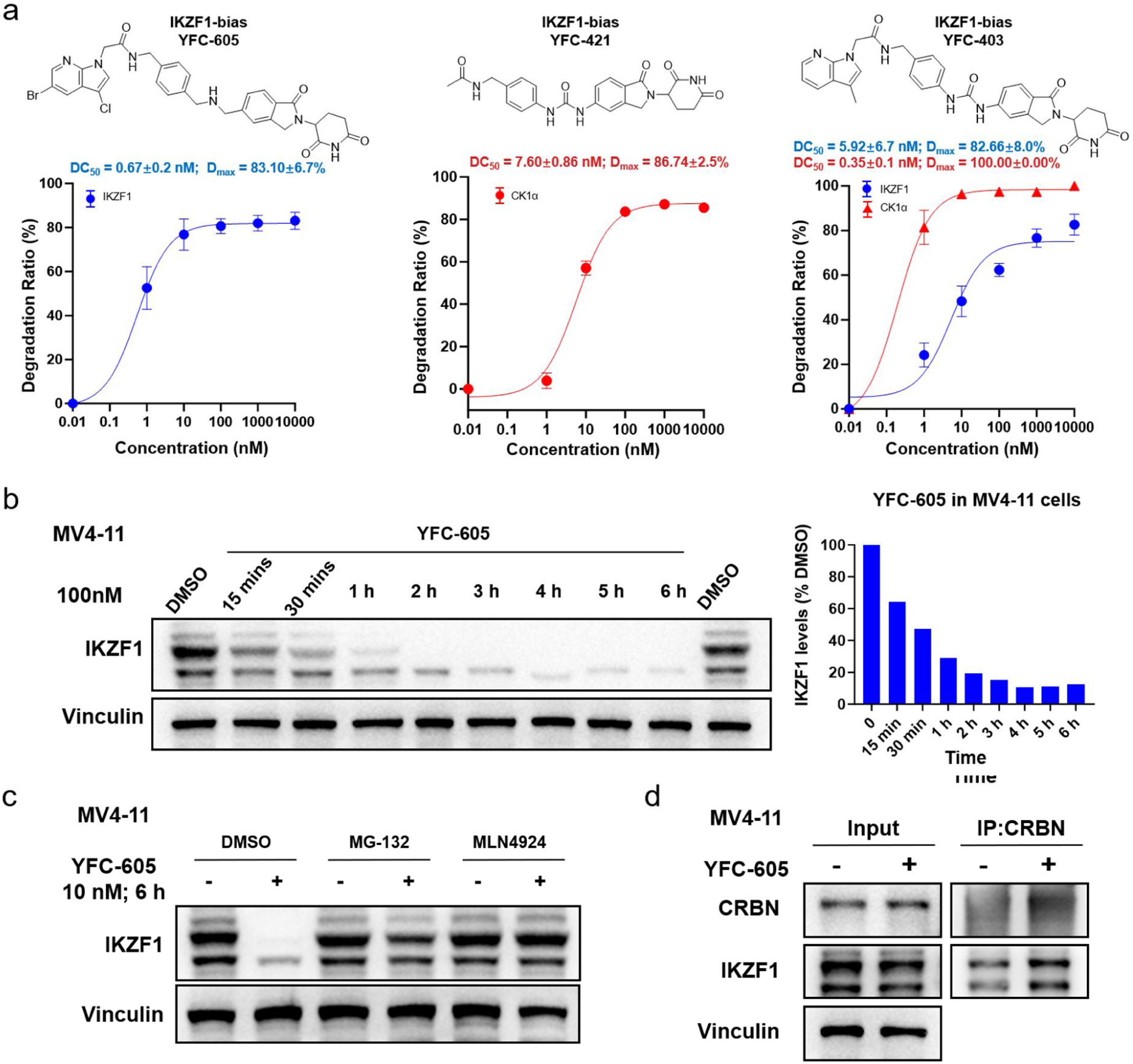
Prospective validation and mechanistic characterization of prioritized CRBN-based molecular glue degraders. **a**, Dose-dependent degradation of IKZF1 and CK1α by representative compounds YFC-605, YFC-421, and YFC-403. **b**, Time-dependent degradation of IKZF1 triggered by 100 nM YFC-605. Vinculin was used as the loading control. **c**, Rescue assays with proteasome inhibitor MG-132 and NAE inhibitor MLN4924, verifying that YFC-605-mediated IKZF1 degradation relies on the proteasome and neddylation pathway. **d**, Co-immunoprecipitation assay was performed to investigate the regulation of CRBN-IKZF1 interaction by YFC-605.

To dissect the degradation mechanism, YFC-605 was selected as a chemical tool given its potent IKZF1 activity. Time-course profiling in MV4-11 cells showed that 100 nM YFC-605 induced rapid and near-complete loss of IKZF1, with substantial depletion apparent within 30 min and maximal degradation sustained through 6 h (Figure 6b), consistent with an event-driven, catalytic mode of action rather than stoichiometric target engagement. To establish pathway dependence, we performed rescue experiments: co-treatment with the proteasome inhibitor MG-132 or the NEDD8-activating enzyme (NAE) inhibitor MLN4924 abolished YFC-605-induced IKZF1 degradation (Figure 6c), indicating that the loss of IKZF1 requires both proteasomal activity and cullin-RING ligase neddylation, the hallmarks of CRL4^CRBN^-mediated turnover. Finally, co-immunoprecipitation showed that YFC-605 markedly enhanced the CRBN-IKZF1 interaction (Figure 6d), demonstrating that the compound acts as a molecular glue that promotes ternary complex formation between CRBN and the IKZF1 neosubstrate. Together, these results confirm that YFC-605 degrades IKZF1 through the canonical molecular-glue mechanism: ligand-induced CRBN engagement, proteasome- and neddylation-dependent ubiquitination, and enhanced CRBN-neosubstrate proximity.

### MG2Act-driven hierarchical prioritization identifies the CDK4 degrader SWC-202

A demanding practical test of a mechanism-aware model is whether it can adapt to targets that are under-represented during pretraining. CDK4 and its close paralog CDK6 are cyclin-dependent kinases that phosphorylate the retinoblastoma protein and drive G1-phase cell-cycle progression, and their aberrant activation is a hallmark of multiple human cancers^37^. Three CDK4/6 inhibitors (palbociclib, ribociclib and abemaciclib) are approved for hormone receptor-positive, HER2-negative breast cancer^41^, but acquired resistance and the inability to achieve sustained kinase-independent suppression of CDK4 function motivate the development of CDK4 degraders. Most CDK4/6 degraders reported to date are bifunctional PROTACs derived from approved CDK4/6 inhibitors, typically tethered to CRBN or VHL ligands^42^. CDK4 contains a CRBN-compatible G-loop degron, and proteomic studies of the CRBN target space have suggested that CDK4 can be engaged as a CRBN-dependent degradable substrate^3^. CDK4 is therefore both biologically motivated and a stringent benchmark for transferability.

To prospectively evaluate whether MG2Act can generalize to target proteins with distinct degron contexts, we deployed the model to screen an in-house CRBN-biased chemical library (n = 167) for novel CDK4 molecular glue degraders. In this workflow, MG2Act served as the core target-conditioned predictive engine, while scaffold-level decomposition and heuristic sampling structured and de-replicated the prioritized candidate pool for chemical synthesis and biological evaluation (Figure 7a).

**Figure 7.**
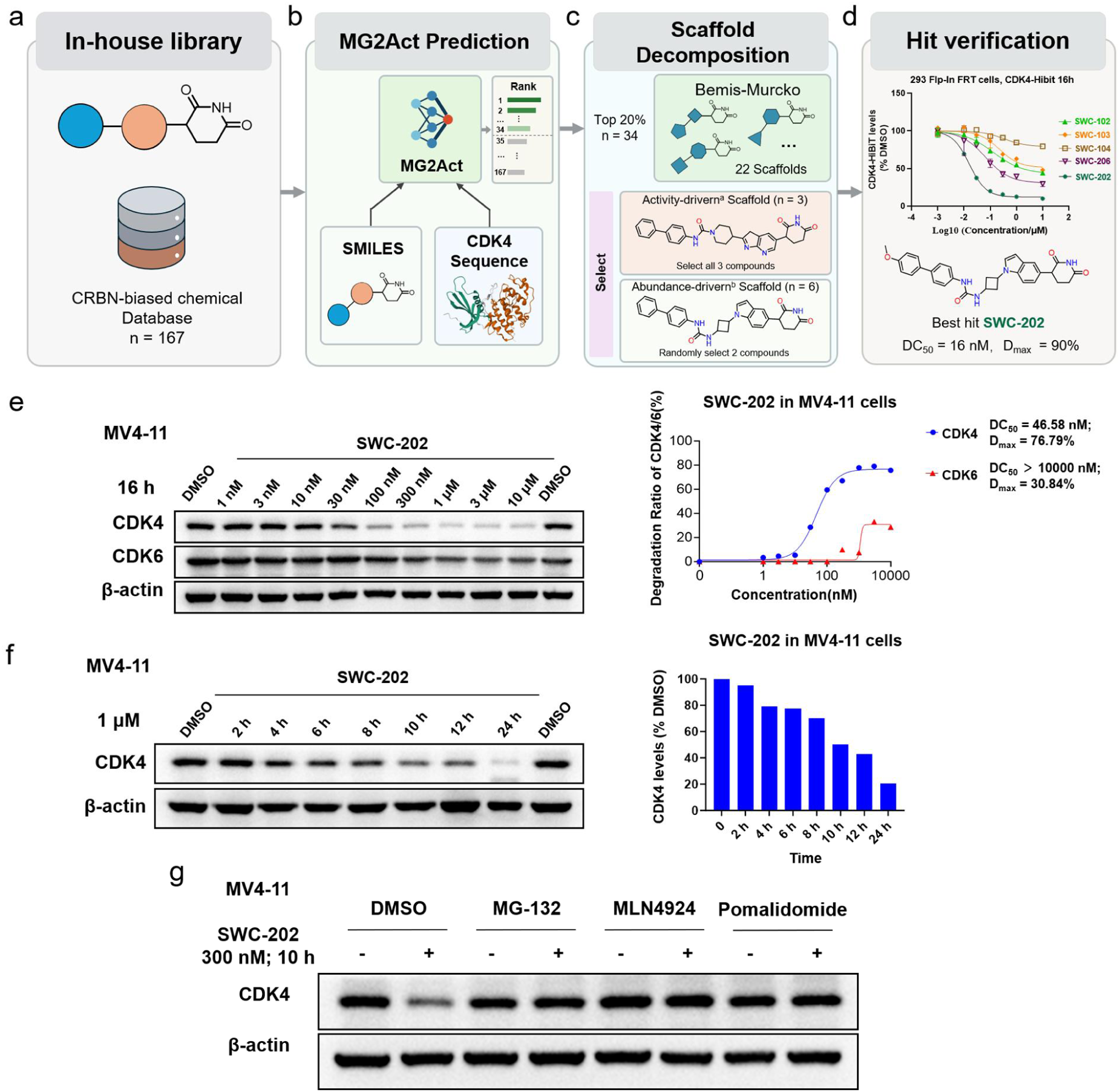
Prospective discovery of a non-classical IMiD-core CDK4 molecular glue (SWC-202) using MG2Act. **a**, In-house CRBN-biased chemical library consisting of 167 glutarimide-containing candidate molecules designed for E3 ligase engagement. **b**, Target-conditioned MG2Act scoring pipeline. Small-molecule SMILES and the human CDK4 protein sequence were integrated into MG2Act to compute predicted degradation activity scores (Score_pred) and rank the 167 candidates. **c**, Bemis– Murcko scaffold decomposition and dual-criterion candidate sampling. The top 20% high-scoring molecules (n = 34) were decomposed into 22 distinct scaffolds. Candidate selection was guided by two complementary criteria: ^a^Activity-driven scaffold (highest mean MG2Act score; all 3 compounds selected), ^b^Abundance-driven scaffold (most populated scaffold, n = 6; 2 representative compounds sampled). **d**, Hit verification and dose–response characterization. SWC-202 achieves DC_50_ = 15.9 nM and D_max_ > 90% in the CDK4-HiBiT assay. **e**, Dose-dependent degradation of CDK4 and CDK6 upon 16 h treatment of SWC-202. β-Actin served as the loading control. Quantification curves for CDK4 and CDK6 degradation are presented on the right. **f**, Time-dependent degradation of CDK4 induced by 1 μM SWC-202. CDK4 abundance was examined via immunoblotting, with β-Actin as the loading control. **g**, SWC-202-induced CDK4 degradation (300 nM, 10 h) is rescued by MG-132 and MLN4924 and competed by pomalidomide, confirming a CRBN-, neddylation- and proteasome-dependent mechanism.

All 167 candidate molecules were first evaluated by MG2Act, combining small-molecule SMILES representations and the CDK4 protein sequence to generate target-conditioned degradation activity scores (Score_pred_) (Figure **7a**). Candidates were ranked based on Score_pred_, and the top 20% (n = 34) high-scoring compounds were prioritized for downstream scaffold-level diversity analysis (Figure **7b**). This primary MG2Act scoring step effectively eliminated low-confidence chemical space while retaining molecules with high predicted target-specific activity.

Subsequently, Bemis–Murcko scaffold decomposition was performed on the 34 prioritized molecules using RDKit to group them into distinct scaffold clusters. Target clusters were isolated using a dual-criterion selection strategy guided directly by MG2Act activity predictions alongside scaffold abundance:

Activity-driven cluster: Selected based on its superior average MG2Act predicted score (Mean Prediction = 0.574), representing the chemotype predicted to exhibit the highest degradation efficacy. For the scaffold possessing the highest average MG2Act score, all associated molecules (n = 3) were retained to ensure the capture of top-predicted active hits. Abundance-driven cluster: Selected based on structural enrichment (n = 6 molecules), representing the dominant chemotype within the top 20% candidate pool. For this most populated scaffold (n = 6), two representative compounds (n = 2) were sampled to capture intra-scaffold structural variations while avoiding redundant testing (Figure **7c**).

Biological evaluation of the prioritized candidates using a CDK4-HiBiT cellular degradation assay confirmed the predictive utility of MG2Act, identifying SWC-202 as a highly potent CDK4 degrader with a DC_50_ value of 15.9 nM and a D_max_ > 90% (Figure **7d**). Notably, SWC-202 features a non-classical IMiD-core framework distinct from canonical glutarimide or phthalimide scaffolds. The successful discovery of SWC-202 demonstrates that MG2Act’s mechanism-inspired architecture effectively captures target-specific compatibility rules. Rather than relying on simple ligand-level similarity, MG2Act acts as a robust predictive engine that, when combined with scaffold-level filtering, efficiently navigates chemical space to prioritize structurally novel and highly active molecular glue degraders.

To further characterize SWC-202 and confirm degradation of the endogenous target, we examined its activity in MV4-11 cells by immunoblotting. Dose-dependent treatment for 16 h produced potent depletion of endogenous CDK4 (DC_50_ = 46.58 nM, D_max_ = 76.79%), while the closely related paralog CDK6 was largely spared (DC_50_ > 10 µM, D_max_ = 30.84%) (Fig. 7e). The higher DC_50_ in this endogenous-protein assay relative to the CDK4-HiBiT reporter system (15.9 nM) is consistent with the differing sensitivities of the two readouts, and the pronounced CDK4-over-CDK6 selectivity indicates that SWC-202 discriminates between paralogous kinases that share high sequence identity but differ in their CRBN-compatible degron context. Time-course analysis showed that 1 µM SWC-202 induced progressive loss of CDK4, with degradation becoming evident by 4–6 h and reaching near-complete depletion by 24 h (Fig. 7f). Rescue experiments further confirmed the mechanism: CDK4 degradation was abolished by the proteasome inhibitor MG-132 and the NAE inhibitor MLN4924, and competed away by the CRBN ligand pomalidomide, establishing that SWC-202 acts through the canonical CRBN binding pocket in a neddylation- and proteasome-dependent manner (Fig. 7g). Together, these cellular data confirm that the MG2Act-prioritized hit SWC-202 is a potent and CDK4-selective degrader of the endogenous kinase, validating the transferability of the model’s directed compatibility signal to a target under-represented during pretraining.

## Discussion

Molecular glue degraders pose a general modelling problem in which activity emerges from directional, cooperative molecular assembly rather than from ligand binding alone. We used CRBN-mediated degradation as a representative, data-rich case to test whether this directionality can be encoded as an architectural inductive bias. CRBN is an ideal starting point because its ligand-induced surface remodeling and neo-substrate recruitment are structurally well established, its available data are more extensive than for other E3 systems, and many existing AI frameworks do not explicitly represent the directed order of molecular glue action.

Three convergent lines of evidence support a mechanism-consistent interpretation. First, reversing the order of the two cross-attention stages, an intervention that targets the directionality of the architecture while keeping other elements fixed, reduces continuous-score regression performance from R² = 0.595 to 0.4833. This R² should be interpreted as a measure of continuous activity ranking rather than binary active or inactive classification. Second, under a stringent de-redundancy benchmark that closes the path of label memorization, baselines decline markedly (R² ≤ 0.2710) while MG2Act retains R² = 0.4223, indicating that its predictive signal arises from learned compatibility rather than rote association alone. Third, a single chemical modification known to abolish CRBN engagement (*N*-methylation of the glutarimide) drives the predicted activity from 0.4396 to 0.1036, and computational target swapping shifts predictions coherently with target identity. Each of these tests probes a property that a pure ligand-memorization predictor would have difficulty reproducing, and MG2Act responds to all three in the direction expected from the biology.

These findings highlight the value of encoding temporal biochemical mechanisms as architectural inductive biases in data-constrained biological AI. Unlike data-abundant domains where black-box architectures suffice, targeted protein degradation suffers from scarce quantitative data, making symmetric fusion assumptions a performance bottleneck. Extending the tradition of domain-informed AI (e.g., PINNs^26^ and P-NET^27^), MG2Act operationalizes the sequential order of biochemical events. This design logic is readily extendable to other E3-first systems (e.g., VHL, DCAF15) and non-degradative proximity-inducing glues wherever molecular assembly follows a defined directional sequence^1^.

Several limitations remain. Directional ablation alone cannot prove that the model has learned the biological mechanism; the supporting interpretation rests on its convergence with chemical perturbation, target swapping, de-redundancy testing and prospective validation. The structure-independent design is an asset for scalability and a deliberate choice given the unreliability of co-folding methods for transient ternary interfaces^18^, but integration of dynamic conformational ensembles from advanced MD or foundation-model-based structure prediction could further refine resolution on context-dependent interaction patterns. The modest absolute correlations against IKZF1 and CK1α in the prospective 28-molecule set (Pearson r = 0.660 and 0.595) reflect the challenge of continuous activity ranking in sparse prospective settings, where hit enrichment and prioritization efficiency are more relevant practical metrics than absolute correlation. Finally, the continuous activity score is currently a hand-designed combination of DC_50_ and D_max_; future work could explore learning this combination end-to-end.

The observed performance drop on out-of-distribution (OOD) targets highlights a general bottleneck in biological machine learning under data-constrained regimes. To explore the model’s extrapolative potential without target-specific retraining, we evaluated MG2Act on CDK4—a target absent from the pre-training dataset. By coupling target-conditioned MG2Act scoring with a scaffold-level hierarchical sampling strategy, we successfully prioritized and validated the non-classical IMiD degrader SWC-202. This provides encouraging initial evidence that encoding the temporal biochemical mechanism as an architectural bias may help cushion data constraints and support candidate prioritization even in sparsely characterized target contexts.

Nevertheless, the predictive scope of the current framework remains inherently bounded by the relatively modest volume and target diversity of available molecular glue data. As high-throughput degradation profiling expands and public repositories grow, incorporating larger, multi-E3 training datasets is expected to further refine MG2Act’s fine-grained activity ranking and broaden its out-of-distribution generalization.

In summary, MG2Act provides a principled, quantitative benchmark for molecular glue modelling. By using CRBN-mediated degradation as a representative E3-first system, it shows how a directional molecular mechanism can be translated into a sequential attention architecture for target-conditioned activity ranking. As an open-access platform for MGD efficacy scoring, MG2Act should be useful for medicinal chemists and computational biologists working on targeted protein degraders, and the design principle it embodies may find applications across degradative and non-degradative molecular glues wherever biology imposes an ordered flow of molecular information.

## Methods

### Chemistry

Detailed synthetic procedures and full chemical characterization (¹H/¹³C NMR, UPLC-MS) of YFC-403, YFC-421, YFC-605, SWC-202 and all other compounds reported in this study will be provided in the Supplementary Information upon peer-reviewed publication.

### Data curation and continuous activity labelling

CRBN-related entries were re-examined from MolGlueDB^28^ and the original literature. Ambiguous or heterogeneous activity readouts were resolved by pharmacology-trained curators, and equivalent values were inferred when explicit DC_50_ measurements were unavailable. DC_50_ values were mapped to a normalized potency score on a 0–1 scale (>10 µM → 0; 1–10 µM → 0.2; 0.1–1 µM → 0.4; 10–100 nM → 0.6; 1–10 nM → 0.8; <1 nM → 1.0). When available, D_max_ was incorporated as a penalty term (>80% → 0; 60–80% → 0.33; 40–60% → 0.67; <40% → 1.0) to account for degradation efficacy beyond potency. The final activity score is a multiplicative combination of normalized DC_50_ and a D_max_-associated penalty:

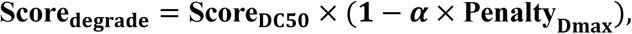

with α a tunable weight (the value used in training is in the Supplementary Information). For entries lacking D_max_, the penalty contribution was minimized. From the initial pool of 1,207 high-confidence compound-target pairs, we retained 21 targets with at least 10 entries each, yielding a refined modeling dataset of 1,159 pairs. To ensure each target was adequately evaluated, the data was distributed via a stratified 8:1:1 split into training, validation, and test sets, maintaining target representation across all three subsets.

### Protein representation

The E3 ligase and the target protein were encoded by the ESM-C 300M protein language model, producing residue-level embeddings:

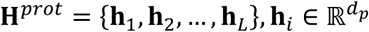

Embeddings were projected into a shared latent space by a learnable linear transformation, with padding masks accommodating variable-length sequences. A global protein representation was obtained by pooling residue-level embeddings along the sequence dimension. The E3 representation used the CRBN functional domain rather than the full sequence (Table 2 ablation).

### Molecular graph and functional-group representation

Each small molecule was parsed by RDKit (v2025.03.1) into an rdkit.Chem.Mol object and represented as a graph G = (V, E), with atoms as nodes and bonds as edges. Atom-level features were extracted from RDKit atom properties (elemental type, degree, aromaticity, hydrogen count, hybridization, formal charge). A three-layer GAT produced atom embeddings:

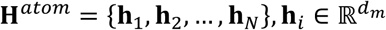

Functional groups were identified by SMARTS-based substructure matching. Starting from 29 commonly occurring groups, we retained those appearing in >5% of molecules and merged chemically related variants, yielding 14 representative functional groups. For each functional group f_k, its embedding is computed by pooling the embeddings of its constituent atoms:

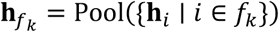

where Pool denotes mean pooling. This results in a compact and interpretable set of functional group representations:

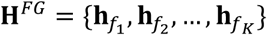

### Functional-group-aware sequential cross-attention

MG2Act applies functional-group-aware cross-attention to model protein-molecule interactions without explicit 3D structures. In each attention head, functional-group embeddings serve as queries while protein residue embeddings serve as keys and values:

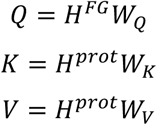

*W_Q_*, *Q_K_* and *W_V_* are learnable projection matrices.

The protein-conditioned functional group representations are computed via scaled dot-product attention:

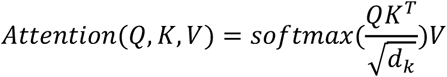

Attention fusion is performed in two sequential stages to reflect the directional biology of CRBN-mediated ternary complex formation. In stage 1, functional groups attend to the E3 ligase (CRBN), yielding E3-conditioned molecular representations that operationalize ligand-induced surface remodelling. In stage 2, these CRBN-conditioned representations attend to the target protein, producing target-conditioned functional-group features.

### Functional-group boosting

To incorporate weak chemical priors derived from experimentally validated molecular glue scaffolds, we introduce a functional group boosting strategy.

For a predefined set of functionally relevant groups F_prior_, their attention weights are softly amplified:

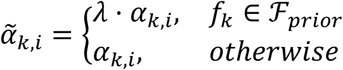

The optimal λ = 2 was selected from validation-set performance (Table 3); the operation can be disabled for ablation studies.

### Multimodal fusion and prediction head

After functional-group–aware attention fusion, protein–molecule interactions have already been explicitly modeled. To integrate complementary information from different modalities, we construct a multimodal representation by aggregating attention-enhanced molecular features and global protein embeddings. Specifically, the protein representations of the E3 ligase and the target protein are obtained by pooling their residue-level embeddings:

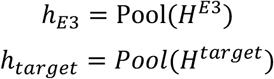

The molecular representation is obtained by pooling the protein-conditioned functional group embeddings:

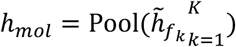

The final multimodal feature vector is constructed by concatenation:

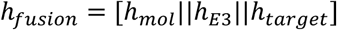

The fused representation is passed through a multilayer perceptron (MLP) to predict molecular glue activity:

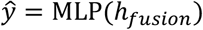

This design decouples interaction modeling, handled by attention-based fusion, from activity prediction, enabling stable optimization and interpretable molecular feature attribution.

### Machine Learning Baseline Models

To rigorously benchmark the performance of MG2Act, three widely used machine learning regression algorithms—Random Forest (RF), Support Vector Regression (SVR), and XGBoost—were implemented using the scikit-learn and xgboost Python libraries. To ensure a strict and fair comparison, all baseline models incorporated identical multimodal information to MG2Act via a target-aware feature concatenation strategy. Each target-aware baseline used a concatenated feature vector comprising the molecular fingerprint, pooled ESM-C embedding of the CRBN CTD, and pooled ESM-C embedding of the corresponding target protein.

### HiBiT degradation assay

The CDK4-HiBiT plasmid vector was obtained from MiaoLingBio (China). CDK4-HiBiT cDNA was cloned into the pcDNA™5/FRT vector and integrated into the genomic FRT site of host cells using the Flp-In™ system (Invitrogen). Stable cell lines were selected with hygromycin B (Sigma, 400051). Integration was verified using the Nano-Glo® HiBiT Lytic Detection System (Envision plate reader, Tecan). CDK4-HiBiT cells were treated with the indicated compounds at concentrations from 0.01 nM to 10 µM or with DMSO as vehicle control for 16 h. The HiBiT luminescent signal was detected using the Nano-Glo® HiBiT Lytic Detection System and dose-response data were fitted to a sigmoidal curve in GraphPad Prism.

### Co-folding prediction and correlation analysis

For each molecular glue–CRBN CTD–target combination, ternary-complex structures were independently predicted using AlphaFold3, Boltz-2 and Protenix, with the CRBN C-terminal thalidomide-binding domain, the corresponding target protein and the molecular glue provided as input components. Five predictions were generated for each complex using each workflow, and the highest-ranked prediction was retained according to the method-specific confidence score. Because interface predicted template modeling scores (ipTM) more directly reflect inter-chain assembly and interface confidence, ipTM-related outputs were prespecified as the primary structural metrics. For each target, 7 AlphaFold3, 9 Boltz-2 and 10 Protenix ipTM-related metrics were included in the analysis. Pearson correlations between each metric and the experimentally measured degradation activity score were calculated separately for CK1α and IKZF1, with MG2Act evaluated on the same compound–target sets. For visualization, model-derived metrics and experimental activity scores were independently z-standardized within each target dataset.

### Cell lines and cell culture

WSU-DLCL2 cells (Wuhan Procell Life Technology) were cultured in RPMI 1640 supplemented with 10% FBS and 1% antibiotic-antimycotic (Procell). HEK293 cells containing the FRT recombination site were provided by the YOC Lab, College of Life Sciences, Jilin University, and cultured in DMEM + 10% FBS + 1% P/S. MV-4-11 cells (Procell, CL-0572) were cultured in IMDM + 10% FBS. MM.1S (Cell Bank, Chinese Academy of Sciences) and 22Rv1 (Procell, CL-0004) cells were cultured in RPMI-1640 + 10% FBS. All cells were maintained at 37 °C in a humidified atmosphere with 5% CO_2_ and were regularly tested to confirm absence of mycoplasma. WSU-DLCL2 cells were seeded in 12-well plates (LABSELECT, 11212) at 1.0 × 10^6^ cells per well in medium containing 10% FBS, and treated with the indicated concentrations of compounds for the indicated times.

### Western blotting

Treated cells were lysed in RIPA buffer (Thermo Fisher Scientific, 89901) supplemented with protease-inhibitor cocktail (Roche, 11697498001). Lysates were centrifuged at 12,000 rpm for 15 min at 4 °C; supernatants were separated by 8% SDS-PAGE and blotted to PVDF membranes. Membranes were blocked in 5% non-fat milk in TBST at room temperature for 1 h, incubated with primary antibodies overnight at 4 °C, then incubated with HRP-conjugated secondary antibodies (Goat anti-rabbit IgG-HRP, P03S02; or Goat anti-mouse IgG-HRP, P03S01) for 1 h at room temperature. Signals were visualized with ECL Western blotting reagent (Bio-Rad, 170506). Primary antibodies: IKZF1 (Cell Signaling Technology, 5443S); Vinculin (CST, R13901S); CK1α (CST, 2655S). Raw blot images are provided in the Supplementary Information.

## Data availability

The raw molecular glue data used in this study are available from the MolGlueDB public database (https://www.molgluedb.com/); the scored and normalized continuous-valued dataset is available on GitHub (https://github.com/Zzy522/MG2Act). Raw gel images are included in the Supporting Information. Source data are provided with this paper. Any remaining data or questions can be obtained from the corresponding author.

## Code availability

The source code of MG2Act, related data preparation scripts and the final model weights are available on GitHub (https://github.com/Zzy522/MG2Act).

## Acknowledgments

We gratefully acknowledge the financial support from the Innovative Drug Research and Development – National Science and Technology Major Project, Project 3 (2026ZD1806900), the Qingdao Science and Technology Demonstration Project under the Qingdao Municipal Science and Technology Bureau (25-1-1-sfgc-2-gx), the Taishan Scholars Program of Shandong Province (tstp20250545), and Strategic Priority Research Program of the Chinese Academy of Sciences (XDB1260301).

## Author contributions

Z.Z., X.W., M.Z. and C.Q. conceived and designed the study. Z.Z. and D.T. developed the code, built the network architecture, and trained the deep learning models. Z.Z., L.G., M.H., M.Y. and Z.-Q.Z. collected, standardized, and processed the molecular glue activity benchmark datasets. Y.W., S.Y., L.C. and Q.H. synthesized and characterized the candidate compounds. X.X., L.G., M.H., M.Y., S.M. and Z.L. performed the cell degradation assays, Western blotting, HiBiT detection, and cellular mechanistic experiments. S.F., D.T. and X.-M.X. conducted the ternary complex modeling and molecular dynamics simulations. Z.Z., D.T., X.X., X.W., M.Z. and C.Q. contributed to the analysis and interpretation of the results. Z.Z., D.T., S.F., X.W. and X.X. wrote the manuscript. X.W., M.Z. and C.Q. offered overall supervision and project administration throughout the project. All authors revised and approved the final manuscript.

