## Supplementary Information for "MG2Act: A Mechanism-Inspired Sequential Attention Framework for Molecular Glue Degradation Prediction"

### Content

**Supplementary Table 1. Distribution of the number of different targets and activities in the constructed molecular glue activity dataset.** ........ **S3**

**Supplementary Table 2. Distribution of the proportions of different targets in the constructed molecular glue activity dataset.** ........ **S4**

**Supplementary Table 3. Mean prediction errors of the MG2Act model for different targets on the test set.** ........ **S5**

**Supplementary Table 4. Predictive performance of MG2Act under varying values of α.** ........ **S5**

**Supplementary Table 5. The data completeness/proportion of $D_{max}$ within the dataset.** **S6**

**Supplementary Table 6. Comparison of sample sizes and target counts between the original test set and the de-biased test set (fine-test set)** ........ **S6**

**Supplementary Figure 1. Evaluation of model generalization and protein-conditioned learning.** ........ **S7**

**Supplementary Figure 2. Scores of original molecules versus after glutarimide N-methylation** ........ **S7**

**Supplementary Table 7. Comparison of experimental and predicted results of the MG2Act model for our 28 molecular glues on the experimental datasets for IKZF1 and CK1α.** ........ **S8**

**Supplementary Figure 3. Western blot images for Supplementary Table 7.** ........ **S9**

**Supplementary Table 8. CDK4-HiBiT degradation assay raw luminescence data.** ........ **S18**

**Supplementary Table 9. Virtual screening scores and rankings of in-house CDK4-targeted compounds predicted by MG2Act, alongside their experimentally validated HiBiT degradation activity.** ........ **S18**

**Preparation procedure for final compounds** ........ **S18**

**Supplementary Table 1. Distribution of the number of different targets and activities in the constructed molecular glue activity dataset.**

| ID | Target | $Score_{degrade}$ | | | | | Total |
|---|---|---|---|---|---|---|---|
| | | [0, 0.2) | [0.2, 0.4) | [0.4, 0.6) | [0.6, 0.8) | [0.8, 1.0] | |
| **1** | CK1α | 20 | 38 | 48 | 28 | 42 | 176 |
| **2** | E4F1 | 8 | 1 | 0 | 9 | 0 | 18 |
| **3** | GSPT1 | 45 | 26 | 30 | 67 | 20 | 188 |
| **4** | IKZF1 | 47 | 14 | 17 | 36 | 25 | 139 |
| **5** | IKZF2 | 35 | 13 | 37 | 25 | 23 | 133 |
| **6** | IKZF3 | 22 | 9 | 18 | 17 | 38 | 104 |
| **7** | $IKZF3^{Q147E}$ | 6 | 0 | 4 | 4 | 0 | 14 |
| **8** | NEK7 | 10 | 5 | 1 | 1 | 2 | 19 |
| **9** | PATZ1 | 7 | 3 | 4 | 4 | 0 | 18 |
| **10** | SALL4 | 28 | 9 | 5 | 9 | 3 | 54 |
| **11** | WEE1 | 0 | 1 | 1 | 6 | 5 | 13 |
| **12** | WIZ | 1 | 2 | 3 | 0 | 4 | 10 |
| **13** | ZFP91 | 8 | 2 | 6 | 8 | 0 | 24 |
| **14** | ZKSC5 | 6 | 0 | 5 | 6 | 0 | 17 |
| **15** | ZNF276 | 8 | 5 | 4 | 20 | 0 | 37 |
| **16** | ZNF517 | 7 | 3 | 9 | 13 | 0 | 32 |
| **17** | ZNF582 | 10 | 0 | 7 | 9 | 0 | 26 |
| **18** | ZNF653 | 11 | 3 | 7 | 16 | 0 | 37 |
| **19** | ZNF654 | 7 | 4 | 9 | 14 | 0 | 34 |
| **20** | ZNF787 | 11 | 2 | 11 | 16 | 0 | 40 |
| **21** | ZNF827 | 9 | 3 | 6 | 8 | 0 | 26 |
| | **Total** | 306 | 143 | 232 | 316 | 162 | 1,159 |

Values are compound counts per $Score_{degrade}$ interval.

**Supplementary Table 2. Distribution of the proportions of different targets in the constructed molecular glue activity dataset.**

| ID | Target | $Score_{degrade}$ | | | | | Total |
|---|---|---|---|---|---|---|---|
| | | [0, 0.2) | [0.2, 0.4) | [0.4, 0.6) | [0.6, 0.8) | [0.8, 1.0] | |
| 1 | CK1α | 11.4% | 21.6% | 27.3% | 15.9% | 23.9% | 100% |
| 2 | E4F1 | 44.4% | 5.6% | 0.0% | 50.0% | 0.0% | 100% |
| 3 | GSPT1 | 23.9% | 13.8% | 16.0% | 35.6% | 10.6% | 100% |
| 4 | IKZF1 | 33.8% | 10.1% | 12.2% | 25.9% | 18.0% | 100% |
| 5 | IKZF2 | 26.3% | 9.8% | 27.8% | 18.8% | 17.3% | 100% |
| 6 | IKZF3 | 21.2% | 8.7% | 17.3% | 16.3% | 36.5% | 100% |
| 7 | $IKZF3^{Q147E}$ | 42.9% | 0.0% | 28.6% | 28.6% | 0.0% | 100% |
| 8 | NEK7 | 52.6% | 26.3% | 5.3% | 5.3% | 10.5% | 100% |
| 9 | PATZ1 | 38.9% | 16.7% | 22.2% | 22.2% | 0.0% | 100% |
| 10 | SALL4 | 51.9% | 16.7% | 9.3% | 16.7% | 5.6% | 100% |
| 11 | WEE1 | 0.0% | 7.7% | 7.7% | 46.2% | 38.5% | 100% |
| 12 | WIZ | 10.0% | 20.0% | 30.0% | 0.0% | 40.0% | 100% |
| 13 | ZFP91 | 33.3% | 8.3% | 25.0% | 33.3% | 0.0% | 100% |
| 14 | ZKSC5 | 35.3% | 0.0% | 29.4% | 35.3% | 0.0% | 100% |
| 15 | ZNF276 | 21.6% | 13.5% | 10.8% | 54.1% | 0.0% | 100% |
| 16 | ZNF517 | 21.9% | 9.4% | 28.1% | 40.6% | 0.0% | 100% |
| 17 | ZNF582 | 38.5% | 0.0% | 26.9% | 34.6% | 0.0% | 100% |
| 18 | ZNF653 | 29.7% | 8.1% | 18.9% | 43.2% | 0.0% | 100% |
| 19 | ZNF654 | 20.6% | 11.8% | 26.5% | 41.2% | 0.0% | 100% |
| 20 | ZNF787 | 27.5% | 5.0% | 27.5% | 40.0% | 0.0% | 100% |
| 21 | ZNF827 | 34.6% | 11.5% | 23.1% | 30.8% | 0.0% | 100% |
| | **Total** | 26.4% | 12.3% | 20.0% | 27.3% | 14.0% | 100% |

Values are the row-wise percentage of compounds falling in each $Score_{degrade}$ interval for a given target.

**Supplementary Table 3. Mean prediction errors of the MG2Act model for different targets on the test set.**

| ID | Target | Prediction error of the test set |
|---|---|---|
| **1** | CK1α | 0.17 |
| **2** | E4F1 | 0.05 |
| **3** | GSPT1 | 0.14 |
| **4** | IKZF1 | 0.11 |
| **5** | IKZF2 | 0.12 |
| **6** | IKZF3 | 0.17 |
| **7** | $IKZF3^{Q147E}$ | 0.05 |
| **8** | NEK7 | 0.21 |
| **9** | PATZ1 | 0.12 |
| **10** | SALL4 | 0.13 |
| **11** | WEE1 | 0.02 |
| **12** | WIZ | 0.25 |
| **13** | ZFP91 | 0.04 |
| **14** | ZKSC5 | 0.06 |
| **15** | ZNF276 | 0.09 |
| **16** | ZNF517 | 0.08 |
| **17** | ZNF582 | 0.01 |
| **18** | ZNF653 | 0.07 |
| **19** | ZNF654 | 0.12 |
| **20** | ZNF787 | 0.06 |
| **21** | ZNF827 | 0.20 |

**Supplementary Table 4. Predictive performance of MG2Act under varying values of α.**

| α | MAE | RMSE | $R^2$ |
|---|---|---|---|
| **0.00** | 0.138 | 0.181 | 0.576 |
| **0.25** | 0.128 | 0.175 | 0.607 |
| **0.50** | 0.124 | 0.175 | 0.595 |
| **0.75** | 0.138 | 0.185 | 0.559 |

The parameter α is an empirical coefficient in the comprehensive degradation scoring formula: $Score_{\text{degrade}} = Score_{DC_{50}} \times \left(1 - \alpha \times Penalty_{D_{max}}\right)$, which modulates the penalty weight of $D_{max}$. A value of α indicates that the $D_{max}$ effect is completely omitted, with larger values imposing a heavier penalty. Here, α = 0.5 was selected as a moderate and well-balanced threshold based on empirical domain conventions.

**Supplementary Table 5. The data completeness/proportion of $D_{max}$ within the dataset.**

| $D_{max}$ Availability | Compound Count | Proportion |
|---|---|---|
| **Available** | 445 | 38.4% |
| **Missing** | 714 | 61.6% |
| **Total** | 1,159 | 100% |

For missing samples where $D_{max}$ values were neither reported in the original literature nor inferable from the text, the $D_{max}$ penalty was omitted/excluded during the calculation of the comprehensive degradation score.

**Supplementary Table 6. Comparison of sample sizes and target counts between the original test set and the de-biased test set (fine-test set).**

| ID | Target | All-test | Fine-test |
|---|---|---|---|
| **1** | CK1α | 18 | 12 |
| **2** | E4F1 | 2 | 0 |
| **3** | GSPT1 | 19 | 16 |
| **4** | IKZF1 | 14 | 5 |
| **5** | IKZF2 | 14 | 12 |
| **6** | IKZF3 | 11 | 9 |
| **7** | $IKZF3^{Q147E}$ | 3 | 0 |
| **8** | NEK7 | 2 | 2 |
| **9** | PATZ1 | 2 | 0 |
| **10** | SALL4 | 6 | 5 |
| **11** | WEE1 | 2 | 1 |
| **12** | WIZ | 3 | 2 |
| **13** | ZFP91 | 3 | 0 |
| **14** | ZKSC5 | 2 | 0 |
| **15** | ZNF276 | 4 | 1 |
| **16** | ZNF517 | 4 | 1 |
| **17** | ZNF582 | 3 | 2 |
| **18** | ZNF653 | 4 | 2 |
| **19** | ZNF654 | 4 | 1 |
| **20** | ZNF787 | 4 | 1 |
| **21** | ZNF827 | 3 | 1 |
| **Total** | | 127 | 73 |

The fine-test set was constructed to eliminate potential shortcut learning. Specifically, molecules that appeared in the combined training/validation set with identical activity across different targets were excluded from the test set, whereas multi-target molecules exhibiting activity variance in the training/validation set were strictly retained to evaluate target-specific selectivity.

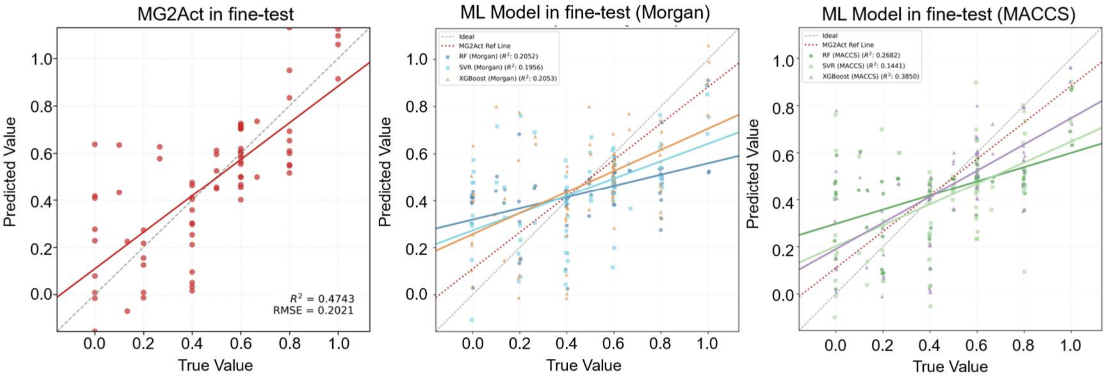


**Supplementary Figure 1. Evaluation of model generalization and protein-conditioned learning.** Comparison of MG2Act and fingerprint-based baselines (RF-ESMC, SVR-ESMC, XGBoost-ESMC) on a stringent test set where training-set-identical molecule–activity associations were excluded. While baseline models show significantly lower performance ($R^2 \leq 0.3850$ for MACCS and $R^2 \leq 0.2053$ for Morgan), MG2Act maintains a robust predictive capacity ($R^2 = 0.4743$). These results indicate that MG2Act's predictive capability reflects a synergistic integration of molecular and protein descriptors, helping to circumvent the shortcut learning pitfalls typical of ligand-only baselines.

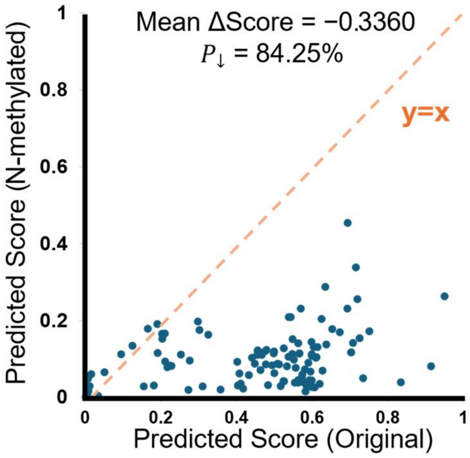


**Supplementary Figure 2. Scores of original molecules versus after glutarimide N-methylation.** A mean reduction of 0.3360 in predicted scores was observed, with 84.25% of molecules showing a predicted decrease, demonstrating strong consistency with the established biological mechanism.

**Supplementary Table 7. Comparison of experimental and predicted results of the MG2Act model for our 28 molecular glues on the experimental datasets for IKZF1 and CK1α.**

| Cpd. | IKZF1 | | | | CK1α | | | |
|---|---|---|---|---|---|---|---|---|
| | $DC_{50}$ (nM) | $D_{max}$ (%) | $Score_{exp}$ | $Score_{pred}$ | $DC_{50}$ (nM) | $D_{max}$ (%) | $Score_{exp}$ | $Score_{pred}$ |
| **YFC-102** | — | — | — | — | >10,000 | 31.11±2.1 | 0 | 0.198 |
| **YFC-106** | — | — | — | — | >10,000 | 46.6±2.9 | 0 | 0.231 |
| **YFC-107** | >10,000 | 18.6±8.6 | 0 | 0.417 | — | — | — | — |
| **YFC-108** | >10,000 | 29.3±5.6 | 0 | 0.4 | — | — | — | — |
| **YFC-201** | — | — | — | — | >10,000 | 30.8±7.3 | 0 | 0.252 |
| **YFC-202** | — | — | — | — | >10,000 | 37.3±12.2 | 0 | 0.236 |
| **YFC-203** | >10,000 | 22.6±6.8 | 0 | 0.408 | — | — | — | — |
| **YFC-204** | >10,000 | 46.19±1.3 | 0 | 0.314 | — | — | — | — |
| **YFC-205** | >10,000 | 35.25±3.7 | 0 | 0.491 | — | — | — | — |
| **YFC-206** | — | — | — | — | >10,000 | 38.95±5.8 | 0 | 0.282 |
| **YFC-207** | >10,000 | 32.84±10.3 | 0 | 0.413 | — | — | — | — |
| **YFC-208** | 33.64 ± 33.7 | 70.27 ± 11.1 | 0.501 | 0.563 | 51.56 ± 12.1 | 65.96 ± 4.8 | 0.501 | 0.38 |
| **YFC-209** | 5.38 ± 1.7 | 60.24 ± 6.7 | 0.668 | 0.445 | >10,000 | 47.40 ± 2.8 | 0 | 0.258 |
| **YFC-210** | 1861.40 ± 1357.3 | 57.76 ± 3.9 | 0.133 | 0.481 | — | — | — | — |
| **YFC-211** | — | — | — | — | >10,000 | 41.57 ± 5.9 | 0 | 0.328 |
| **YFC-212** | 0.257 ± 0.108 | 52.9 ± 2.9 | 0.665 | 0.444 | >10,000 | 41.5 ± 9.7 | 0 | 0.259 |
| **YFC-302** | >10,000 | 40.3 ± 15.3 | 0 | 0.447 | 25.35 ± 16.22 | 62.6 ± 1.3 | 0.501 | 0.341 |
| **YFC-304** | — | — | — | — | >10,000 | 36.6 ± 11 | 0 | 0.256 |
| **YFC-305** | — | — | — | — | 214.97 ± 127.9 | 54.68 ± 3 | 0.266 | 0.221 |
| **YFC-307** | >10,000 | 34.89 ± 8.4 | 0 | 0.484 | >10,000 | 42.59 ± 6.2 | 0 | 0.347 |
| **YFC-403** | 5.92 ± 6.7 | 82.66 ± 8.0 | 0.8 | 0.475 | 0.35 ± 0.1 | 100 ± 0 | 1 | 0.482 |
| **YFC-404** | — | — | — | — | >10,000 | 12.68 ± 6.0 | 0 | 0.336 |
| **YFC-412** | >10,000 | 44.8 ± 5.5 | 0 | 0.19 | >10,000 | 38.9 ± 8.6 | 0 | 0.296 |
| **YFC-415** | >10,000 | 23.1 ± 4.1 | 0 | 0.291 | — | — | — | — |
| **YFC-417** | >10,000 | 19.4 ± 9.1 | 0 | 0.304 | — | — | — | — |
| **YFC-421** | — | — | — | — | 7.6 ± 0.86 | 86.7 ± 2.5 | 0.8 | 0.482 |
| **YFC-604** | 6.66 ± 2.1 | 86.63 ± 6.1 | 0.8 | 0.662 | >10,000 | 6.82 ± 6.5 | 0 | 0.538 |
| **YFC-605** | 0.67 ± 0.2 | 83.10 ± 6.7 | 1 | 0.614 | — | — | — | — |

Degradation was measured by Western blot in WSU-DLCL2 cells after 6 h of compound treatment. "—" indicates the compound was not tested against that target. The experimental degradation score was calculated as:

$$Score_{exp} = Score_{DC_{50}} \times (1 - 0.5 \times Penalty_{D_{max}})$$

where $Score_{DC_{50}}$ was assigned from $DC_{50}$: 1 (<1 nM), 0.8 (1–10 nM), 0.6 (10–100 nM), 0.4 (0.1–1 μM), 0.2 (1–10 μM), and 0 (>10 μM); and $Penalty_{D_{max}}$ was assigned from the maximal degradation efficacy $D_{max}$: 0 (>80%), 0.33 (60–80%), 0.67 (40–60%), 1.0 (<40%). $Score_{pred}$ is

MG2Act predicted score. Across compounds with paired experimental and predicted values, the MG2Act score correlated with the experimental score with Pearson r = 0.660 (IKZF1) and r = 0.595 (CK1α). Data are the mean of at least three biological replicates.

**Supplementary Figure 3. Western blot images for Supplementary Table 7.**

All data are presented as the mean of at least three biological replicates.

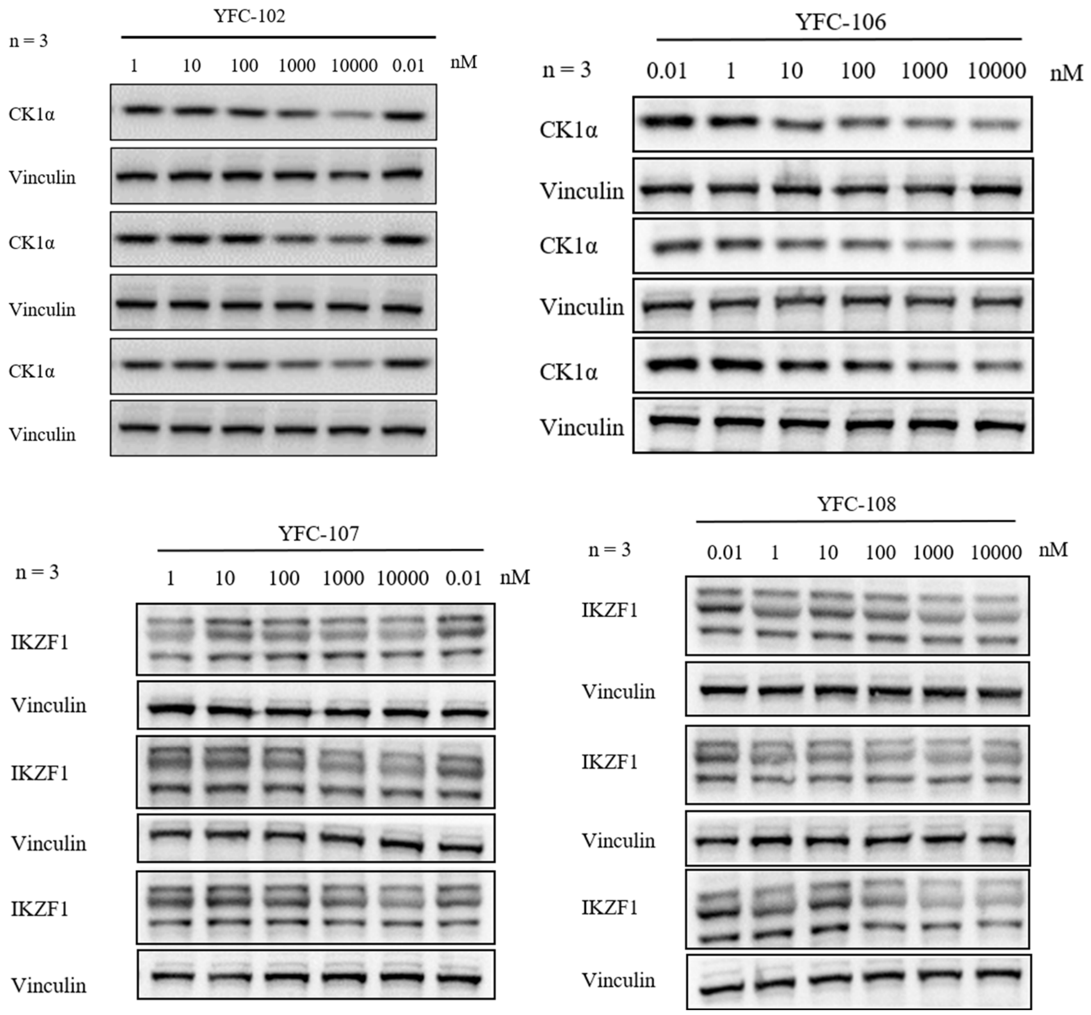

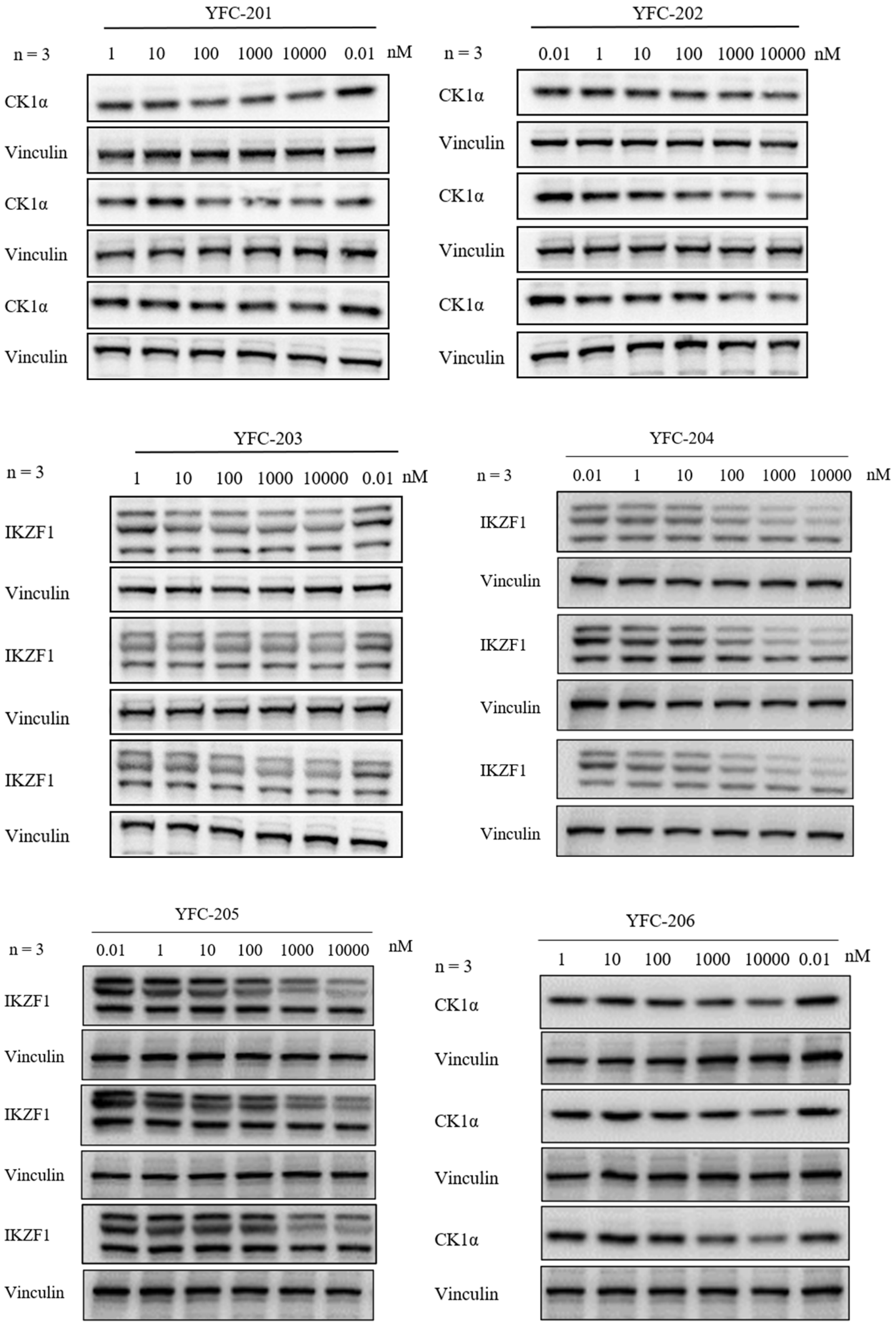
YFC-201
n = 3
1 10 100 1000 10000 0.01 nM
CK1α
Vinculin
CK1α
Vinculin
CK1α
Vinculin
YFC-202
n = 3
0.01 1 10 100 1000 10000 nM
CK1α
Vinculin
CK1α
Vinculin
CK1α
Vinculin
YFC-203
n = 3
1 10 100 1000 10000 0.01 nM
IKZF1
Vinculin
IKZF1
Vinculin
IKZF1
Vinculin
YFC-204
n = 3
0.01 1 10 100 1000 10000 nM
IKZF1
Vinculin
IKZF1
Vinculin
IKZF1
Vinculin
YFC-205
n = 3
0.01 1 10 100 1000 10000 nM
IKZF1
Vinculin
IKZF1
Vinculin
IKZF1
Vinculin
YFC-206
n = 3
1 10 100 1000 10000 0.01 nM
CK1α
Vinculin
CK1α
Vinculin
CK1α
Vinculin

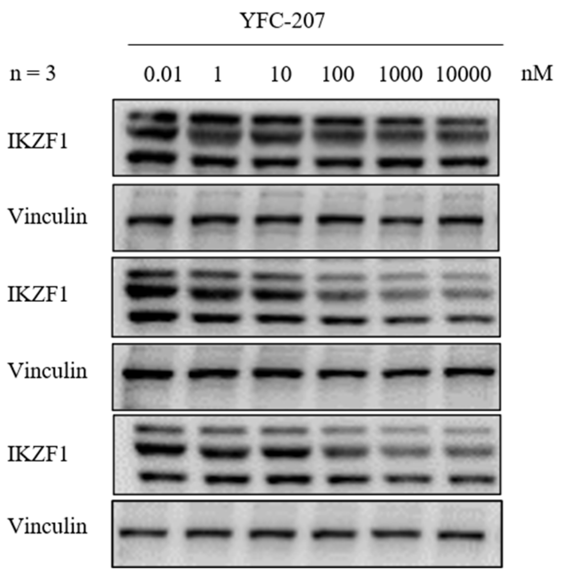
YFC-207
n = 3
0.01 1 10 100 1000 10000 nM
IKZF1
Vinculin
IKZF1
Vinculin
IKZF1
Vinculin

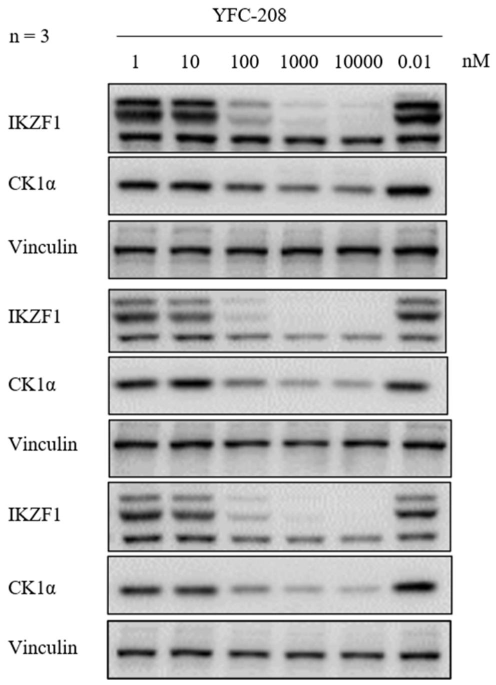
YFC-208
n = 3
1 10 100 1000 10000 0.01 nM
IKZF1
CK1α
Vinculin
IKZF1
CK1α
Vinculin
IKZF1
CK1α
Vinculin

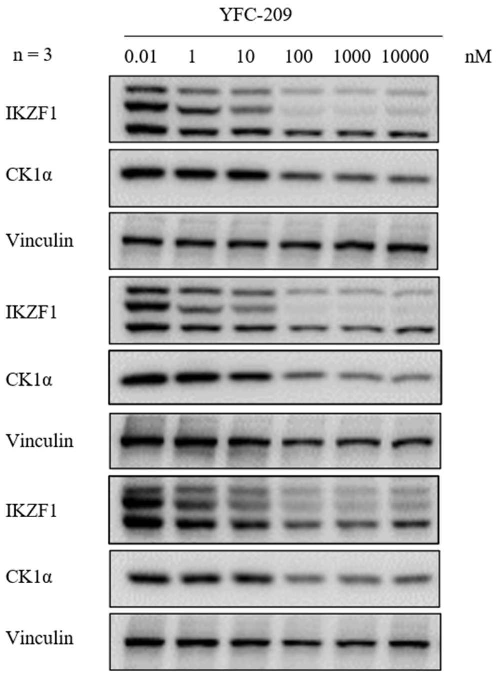
YFC-209
n = 3
0.01 1 10 100 1000 10000 nM
IKZF1
CK1α
Vinculin
IKZF1
CK1α
Vinculin
IKZF1
CK1α
Vinculin

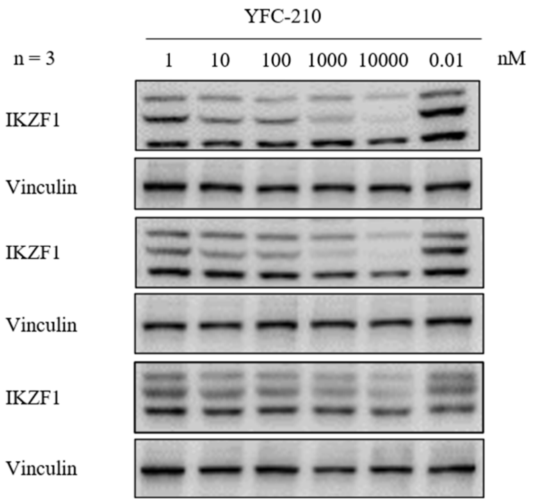
YFC-210
n = 3
1 10 100 1000 10000 0.01 nM
IKZF1
Vinculin
IKZF1
Vinculin
IKZF1
Vinculin

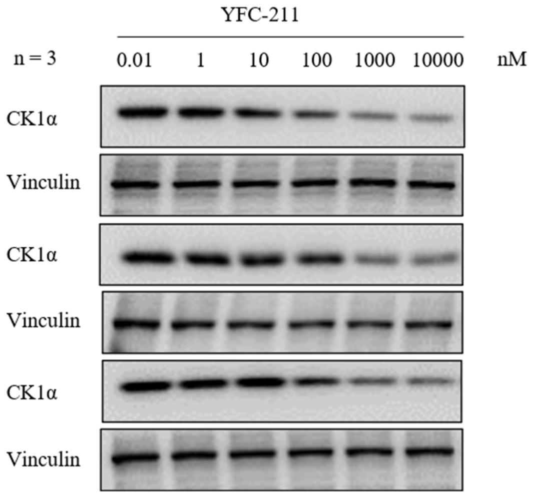
YFC-211
n = 3
0.01 1 10 100 1000 10000 nM
CK1α
Vinculin
CK1α
Vinculin
CK1α
Vinculin

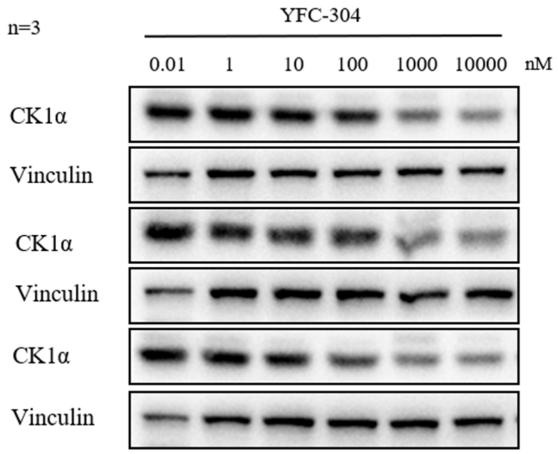
n=3
YFC-304
0.01 1 10 100 1000 10000 nM
CK1α
Vinculin
CK1α
Vinculin
CK1α
Vinculin

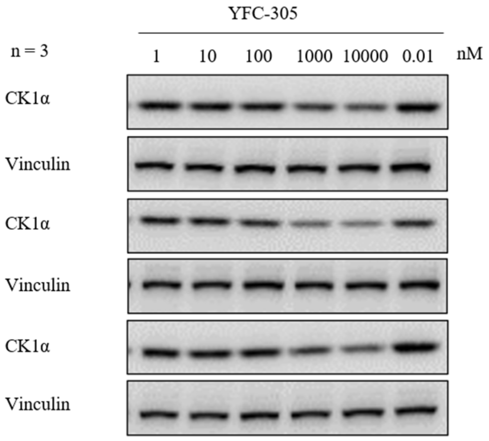
n = 3
YFC-305
1 10 100 1000 10000 0.01 nM
CK1α
Vinculin
CK1α
Vinculin
CK1α
Vinculin

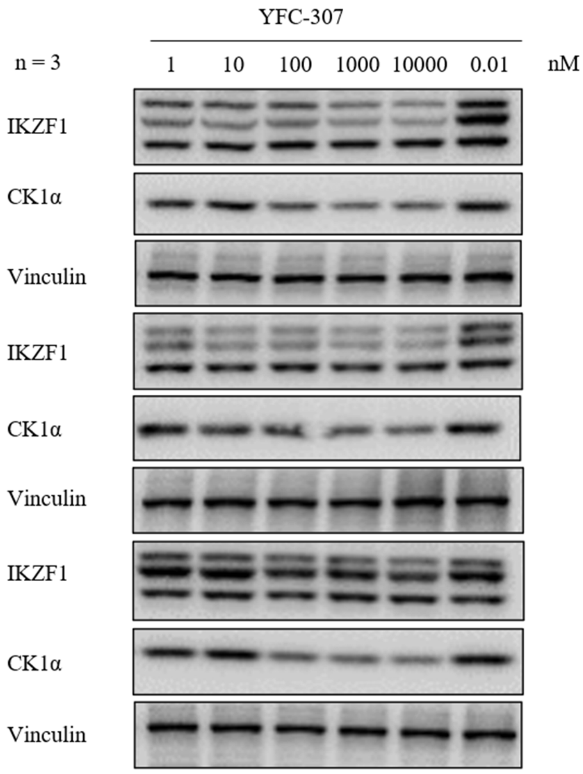
YFC-307
n = 3
1 10 100 1000 10000 0.01 nM
IKZF1
CK1α
Vinculin
IKZF1
CK1α
Vinculin
IKZF1
CK1α
Vinculin

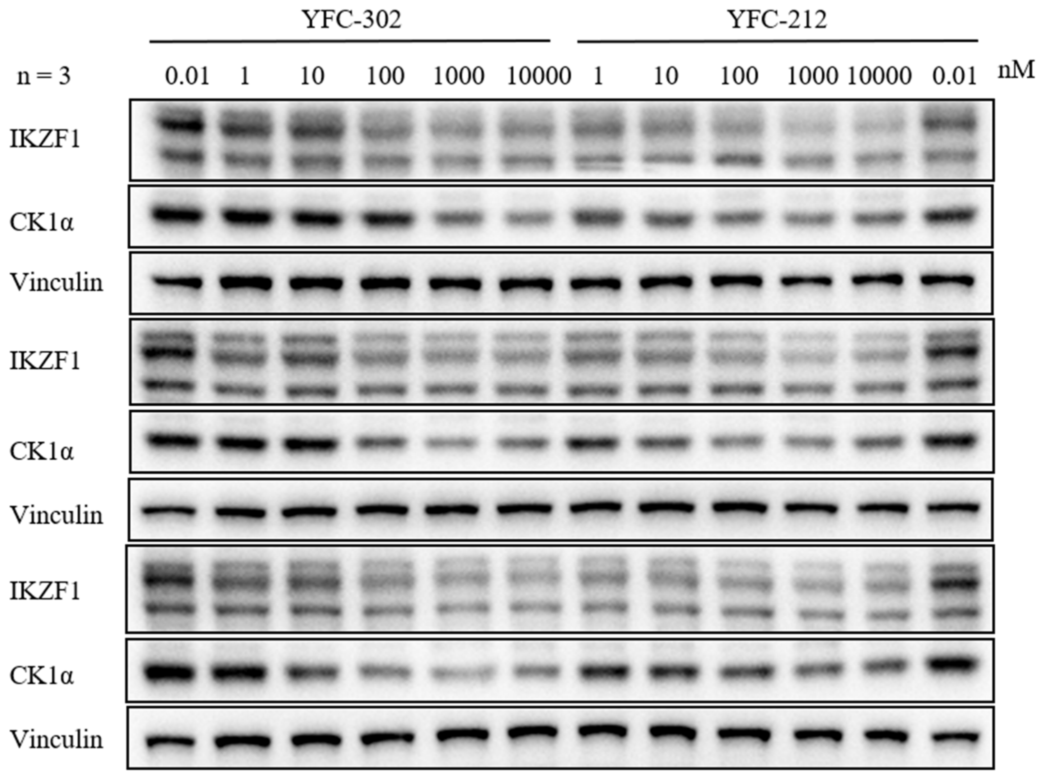
YFC-302
YFC-212
n = 3
0.01 1 10 100 1000 10000 1 10 100 1000 10000 0.01 nM
IKZF1
CK1α
Vinculin
IKZF1
CK1α
Vinculin
IKZF1
CK1α
Vinculin

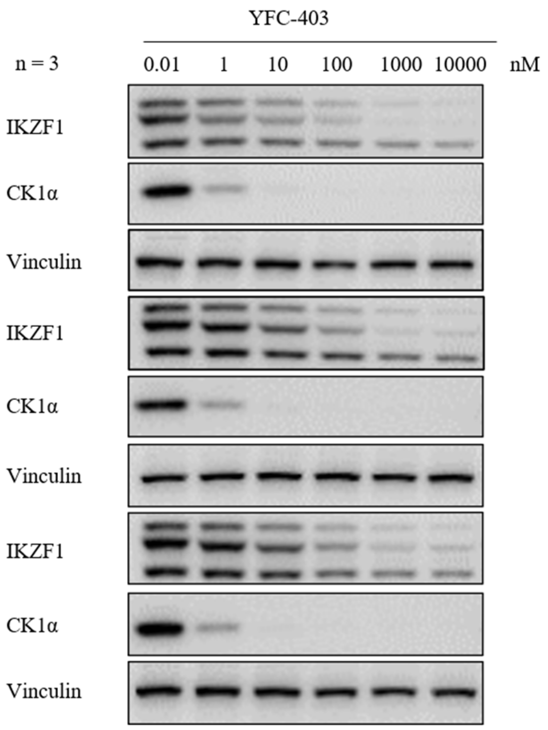
YFC-403
n = 3
0.01 1 10 100 1000 10000 nM
IKZF1
CK1α
Vinculin
IKZF1
CK1α
Vinculin
IKZF1
CK1α
Vinculin

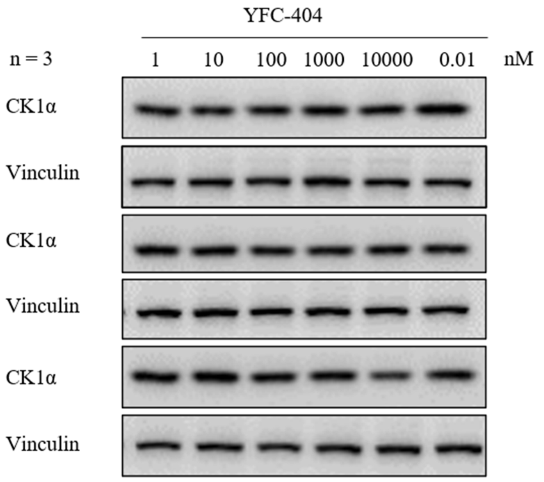
YFC-404
n = 3
1 10 100 1000 10000 0.01 nM
CK1α
Vinculin
CK1α
Vinculin
CK1α
Vinculin

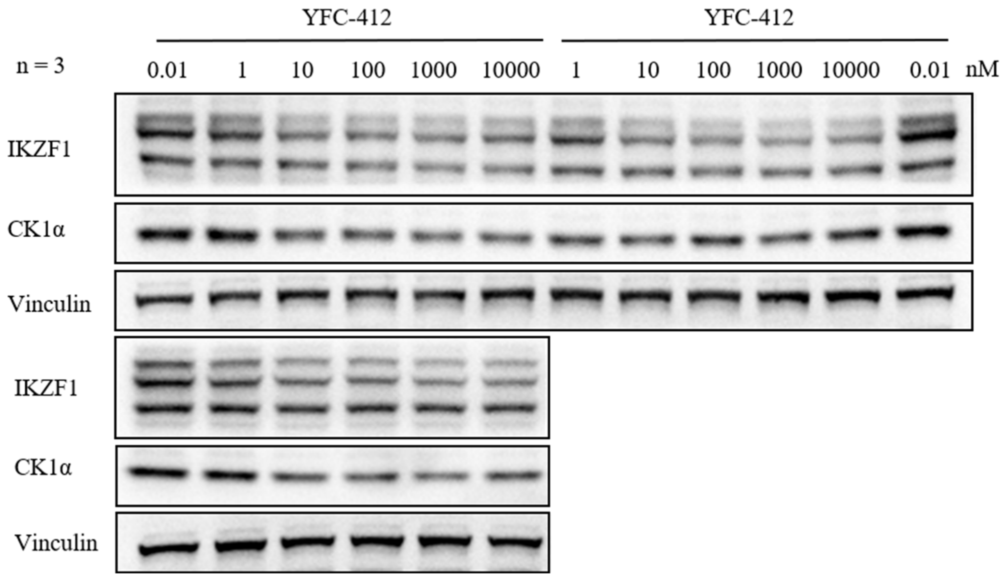
YFC-412
YFC-412
n = 3
0.01 1 10 100 1000 10000 1 10 100 1000 10000 0.01 nM
IKZF1
CK1α
Vinculin
IKZF1
CK1α
Vinculin

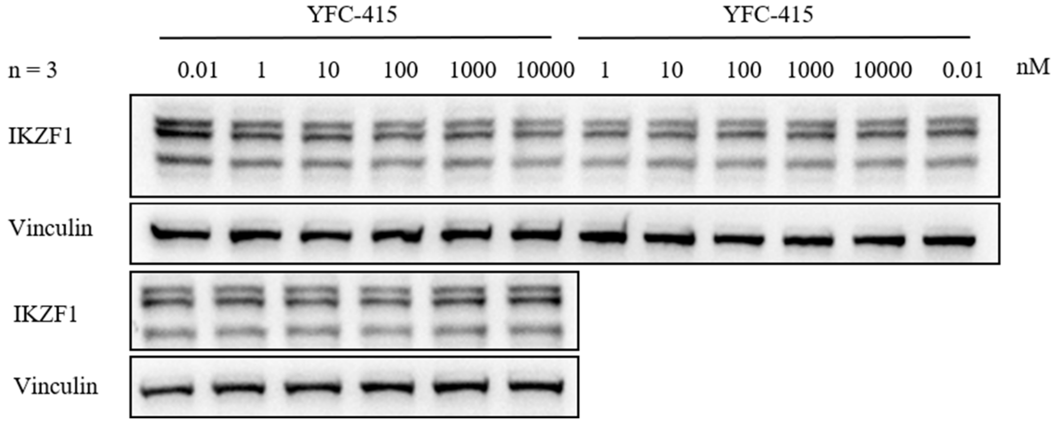
YFC-415
YFC-415
n = 3
0.01 1 10 100 1000 10000 1 10 100 1000 10000 0.01 nM
IKZF1
Vinculin
IKZF1
Vinculin

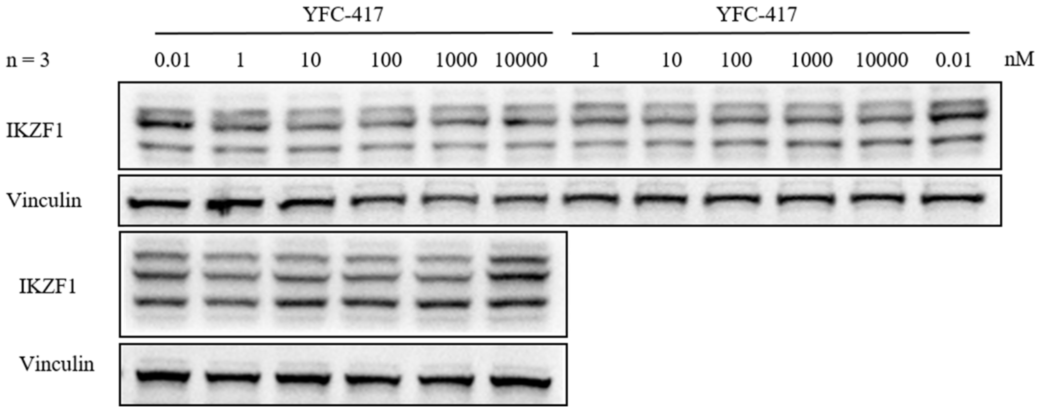
YFC-417
YFC-417
n = 3
0.01 1 10 100 1000 10000 1 10 100 1000 10000 0.01 nM
IKZF1
Vinculin
IKZF1
Vinculin

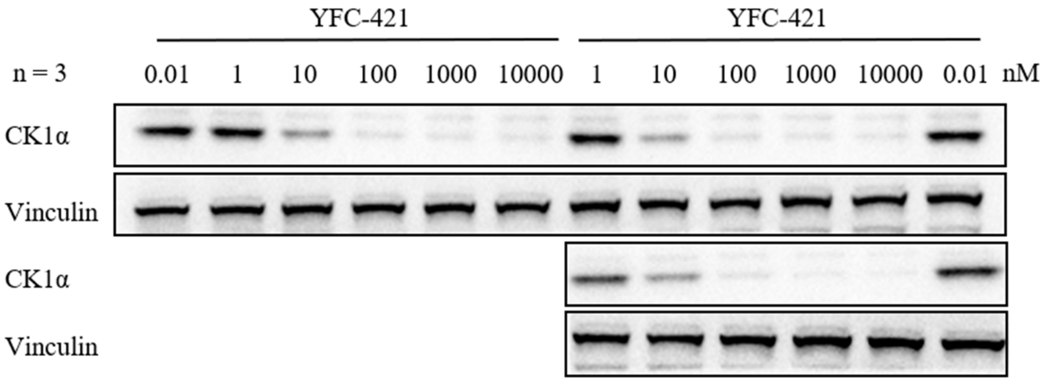

YFC-421
YFC-421
n = 3
0.01 1 10 100 1000 10000 1 10 100 1000 10000 0.01 nM
CK1α
Vinculin
CK1α
Vinculin


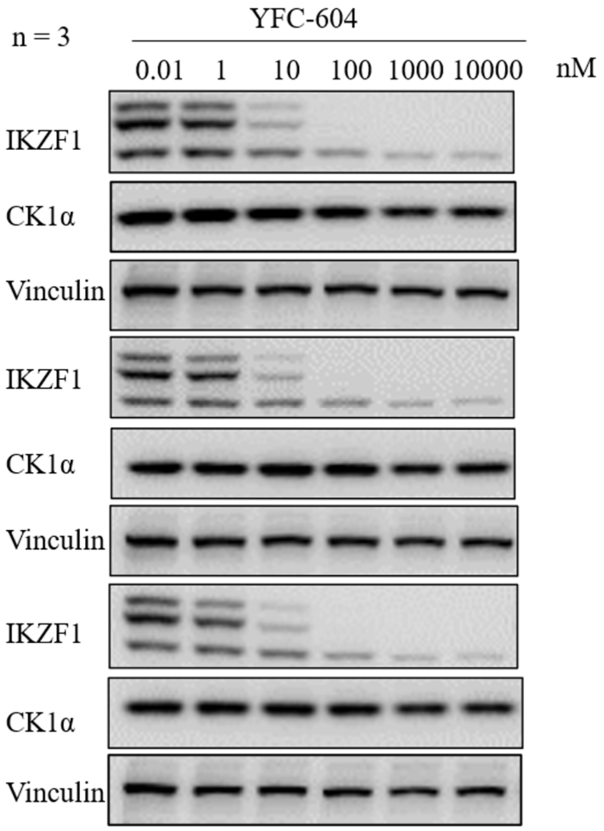

n = 3
YFC-604
0.01 1 10 100 1000 10000 nM
IKZF1
CK1α
Vinculin
IKZF1
CK1α
Vinculin
IKZF1
CK1α
Vinculin

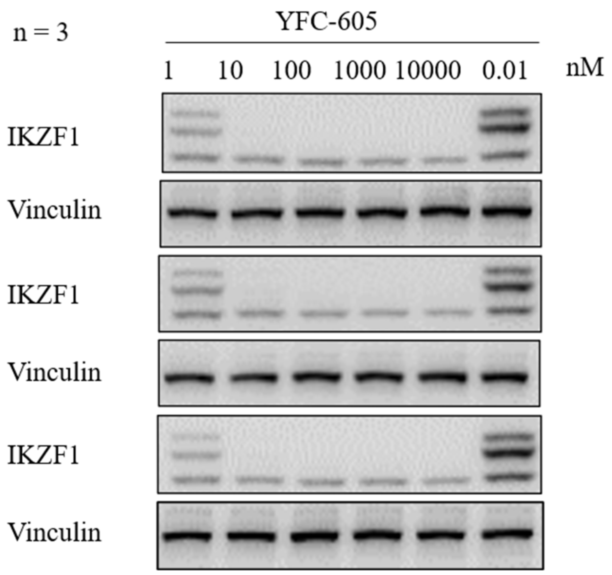
n = 3
YFC-605
1
10
100
1000
10000
0.01
nM
IKZF1
Vinculin
IKZF1
Vinculin
IKZF1
Vinculin

**Supplementary Table 8. CDK4-HiBiT degradation assay raw luminescence data.**

| Cmp. | Rep | CDK4-HiBiT remaining level (% DMSO) at Concentrations (μM, in 293 FRT CDK4-Hibit 16h) | | | | | | |
|---|---|---|---|---|---|---|---|---|
| | | 0 | 10 | 30 | 100 | 300 | 1,000 | 10,000 |
| **SWC-202** | Rep1 | 100.20 | 69.80 | 36.70 | 18.30 | 14.00 | 11.60 | 9.50 |
| | Rep2 | 99.80 | 66.70 | 37.60 | 20.40 | 16.40 | 13.00 | 10.10 |
| **SWC-206** | Rep1 | 99.60 | 92.20 | 85.50 | 64.30 | 47.80 | 39.60 | 33.40 |
| | Rep2 | 100.40 | 97.90 | 90.10 | 59.60 | 45.20 | 46.00 | 30.30 |
| **SWC-102** | Rep1 | 97.30 | 89.90 | 79.90 | 69.40 | 58.70 | 52.60 | 41.10 |
| | Rep2 | 100.30 | 92.50 | 85.90 | 72.30 | 60.60 | 53.20 | 43.60 |
| **SWC-103** | Rep1 | 98.50 | 99.00 | 89.50 | 76.60 | 65.20 | 58.50 | 47.00 |
| | Rep2 | 101.70 | 102.90 | 91.80 | 89.20 | 67.50 | 62.70 | 45.70 |
| **SWC-104** | Rep1 | 99.40 | 96.70 | 96.00 | 88.20 | 87.50 | 81.30 | 76.40 |
| | Rep2 | 102.80 | 97.30 | 99.00 | 90.40 | 87.60 | 80.50 | 76.30 |

CDK4-HiBiT cells were incubated with the indicated concentrations of compounds for 16 h. HiBiT luminescence was detected using the Nano-Glo® HiBiT Lytic Detection System. Values represent individual replicate measurements (Rep1, Rep2) normalized to the DMSO vehicle control (0 nM). These raw data were used for sigmoidal dose-response curve fitting in GraphPad Prism.

**Supplementary Table 9. Virtual screening scores and rankings of in-house CDK4-targeted compounds predicted by MG2Act, alongside their experimentally validated HiBiT degradation activity.**

| ID | MG2Act Prediction | | Measured activity | | |
|---|---|---|---|---|---|
| | rank | Prediction | $Score_{DC50}$[a] | $Penalty_{Dmax}$[b] | $Score_{degrade}$[c] |
| **SWC-102** | 3 | 0.5939 | 0.4 | 0.67 | 0.27 |
| **SWC-103** | 6 | 0.5641 | 0.4 | 0.67 | 0.27 |
| **SWC-104** | 5 | 0.5637 | 0 | 1.00 | 0 |
| **SWC-202** | 14 | 0.4453 | 0.6 | 0 | 0.60 |
| **SWC-206** | 54 | 0.4091 | 0.6 | 0.33 | 0.50 |

[a] $Score_{DC50}$ assignment: 1.0 (<1 nM), 0.8 (1–10 nM), 0.6 (10–100 nM), 0.4 (0.1–1 μM), 0.2 (1–10 μM), 0 (>10 μM).

[b] $Penalty_{Dmax}$ assignment: 0 (>80%), 0.33 (60–80%), 0.67 (40–60%), 1.00 (<40%).

[c] $Score_{degrade} = Score_{DC50} \times (1 - 0.5 \times Penalty_{Dmax})$

#### Preparation procedure for final compounds

**Full synthetic procedures and characterization will be provided upon peer-reviewed publication.**
